# Exploring the sequence and structural determinants of the energy landscape from thermodynamically stable and kinetically trapped subtilisins: ISP1 and SbtE

**DOI:** 10.1101/2024.09.08.611919

**Authors:** Miriam R. Hood, Susan Marqusee

## Abstract

A protein’s energy landscape, all accessible conformations, their populations, and dynamics of interconversion, is encoded in its primary sequence. While how a protein’s primary sequence encodes its native state is well understood, how sequence encodes the kinetic barriers such as unfolding and refolding is not. Here we have looked at two subtiliase homologs from the *Bacillus subtilis*, Intracellular Subtilisin Protease 1 (ISP1) and Subtilisin E (SbtE), that are expected to have very different dynamics. As an intracellular protein, ISP1 has a small pro-domain thought to act simply as a zymogen, whereas the extracellular SbtE has a large pro-domain required for folding. The stability and kinetics of ISP1 and ProSbtE have been previously characterized. We now directly compare their energy landscapes with and without the pro-domain, examining global and local energetics of the mature proteases and the effect of each pro-domain. We find that ISP1’s pro-domain has limited impact on the energy landscape of the mature protein while without the pro-domain, SbtE is thermodynamically unstable and kinetically trapped. The pro-domains show opposite effects on the flexibility of the core of the protein: in the absence of its pro-domain, ISP1’s core becomes more flexible while SbtE’s core becomes more rigid. ISP1 contains a unique conserved insertion, which points to a potential source for these differences. These homologs are an example of how changes in the primary sequence can dramatically alter a proteins energy landscape, and highlight the need for large scale, high-throughput studies on the relationship between primary sequence and conformational dynamics.

## Introduction

Proteins are an essential group of macromolecules, playing important roles in nearly all biological functions. For most proteins, the active conformation is a compact, folded structure whose topology and dynamics are finely tuned and encoded in the linear sequence of amino acids. Protein folding, the process needed to obtain this structure, must take place on a timescale consistent with the protein’s function and environment. Similarly, once folded, both the global and local dynamics must occur on the time- and size-scale needed for function and growth of the organism. The process of protein folding/unfolding occurs along an energy landscape, often referred to as a funnel, which contains all the partially folded states accessible to the protein as well as the kinetic barriers separating them. This entire landscape is encoded in a protein’s amino acid sequence. Despite significant advances in our ability to predict the folded conformation from sequence, (Jumper et al., 2021), we have a relatively poor understanding of how the lifetime or barriers between conformations are encoded. These barriers play numerous, important roles in biology – such as preventing inappropriate proteolytic degradation, as timers for complex systems, and in the pathogenesis of misfolding diseases (Sanchez-Ruiz, 2010). These barriers are not simply a result of the fold or topology of the protein, as proteins with the same overall native fold can have vastly different landscapes (Spudich, Miller, & Marqusee, 2004; Young, Skordalakes, & Marqusee, 2007).

While the native state of most proteins is thought to be at the free energy minimum on this landscape (Anfinsen’s “thermodynamic hypothesis”), such thermodynamic control is not universal. There are now several published examples of proteins for whom the native state is kinetically trapped and the native conformation is maintained by kinetic control (an extremely high barrier to unfolding) (Baker & Agard, 1994a; Sanchez-Ruiz, 2010). In many of these cases, a chaperone assists the protein in getting over the kinetic barrier – often a genetically encoded pro-region of the protein that is cleaved once the protein folds into its native state, leaving a kinetically-trapped native protein. Alpha-lytic protease (αLP), perhaps the most well characterized example of such a protein, was shown to have limited fluctuations from the native state (a high degree of rigidity) and significant resistance to proteolytic degradation (Jaswal, Sohl, Davis, & Agard, 2002a). This led to the widely accepted hypothesis that kinetically trapped, secreted proteases may have evolved such large barriers to improve survival in harsh, protease-rich extracellular environments (P. N. Bryan, 2002; Jaswal et al., 2002a; Sanchez-Ruiz, 2010). The mechanism by which αLP’s large kinetic barrier arose is unknown. It is also unknown whether this is a general feature for all secreted proteases or why a protein would evolve such an unusually large kinetic barrier.

Kinetic barriers are not simply a feature of the topology or conformation, as proteins with similar folds can have dramatically different lifetimes. In fact, proteins with the same topology can show both thermodynamic and kinetic control (Baker & Agard, 1994b; Baker, Sohl, & Agard, 1992a; Sohl, 1997) A deeper understanding of how kinetic barriers are encoded would expand our understanding of how the energy landscape is modulated and allow for the design of conformations with specified dynamics and lifetimes. Here, we investigate an extreme case where two protein sequences from the same organism with the same overall native conformation show dramatic differences in their unfolding barriers. We characterize and compare the energy landscape of both an intracellular and extracellular subtiliase from *Bacillus subtilis*.

Bacterial subtiliases fall into two subfamilies with high structural and sequence conservation: intracellular subtilisin proteases (ISPs) and extracellular subtilisin proteases (ESPs). While these proteins have 30-50% sequence identity and similar folds, paralogs found in the same *Bacillus* species have been shown to have very different folding landscapes (Subbian, Yabuta, & Shinde, 2004; Vévodová et al., 2010). ISPs are thermodynamically stable proteins synthesized with a small (approximately 20 amino acids), N-terminal intramolecular inhibitor – the pro-domain, which is not required for folding (Gamble, Künze, Dodson, Wilson, & Jones, 2011a; Strongin et al., 1978; Subbian et al., 2004). In contrast, ESPs, like other secreted proteases, are at best marginally stable (thermodynamically) and are thought to maintain their fold through a high kinetic barrier to unfolding (P. N. Bryan, 2002; Eder & Fersht, 1995; Sohl, Jaswal, & Agard, 1998; James A Wells & Estell, 1988). ESPs require a 70 to 100 amino acid intramolecular chaperone, the ESP pro-domain, to reach their native state. Once the protein is folded, the pro-domain is auto-digested, trapping the protein in the native state (P. N. Bryan, 2002; P. Bryan et al., 1995; Yabuta, Takagi, Inouye, & Shinde, 2001). While several ESPs (including those studied here) have been characterized biophysically and engineered for industrial purposes, what aspects of their primary sequence convey their impressive kinetic barrier is unclear (Fu, Inouye, & Shinde, 2000; James A Wells & Estell, 1988; You & Arnold, 1996).

Despite the high resource cost of synthesizing a 100 amino-acid chaperone for a single protein-folding event, the ESP pro-domain remains a conserved feature within the subfamily. Sequence analysis shows evidence for positive selection on the pro-domain’s size and charge during the evolution of ISPs, perhaps to facilitate protein degradation and turnover and reduce excessive proteolytic activity in the cell (Subbian et al., 2004). This analysis highlights a handful of surface residues potentially under evolutionary selection, it does not reveal what aspects of sequence or structure account for the differences in folding mechanism. While ISPs have been characterized, what effect the pro-domain may have on their energy landscape has not been fully evaluated. Although, it is assumed that the ISP’s pro-domain’s main function is as an inhibitor and may have minimal impact on the energy landscape (Gamble, Künze, Dodson, Wilson, & Jones, 2011b).

There are many potential environmental pressures that may select for increased unfolding barriers for secreted proteases, including a lack of proteostasis machinery outside the cell, the need to function in a harsh environment, and prevention of autolysis or degradation by competing proteases (Jaswal et al., 2002a; Sanchez-Ruiz, 2010). It is possible that ESPs’ high kinetic barriers, like αLP’s, have evolved to prevent degradation in the extracellular environment; however, this has not been established. Subtilisins are produced by *Bacillus* bacteria which are found largely in the soil but span a wide range of environments from temperate to extreme. In order to survive these inhospitable conditions, *Bacilli* form long-lasting, metabolically-inactive endospores (Turnbull, 1996). Germination of these spores can be triggered by L-amino acids (alanine, valine, and asparagine) – which begins the breakdown of the endospore cortex, allowing cell expansion and water uptake to occur and facilitate the return of metabolism (Peter, 2014). ESPs are believed to degrade other extracellular proteins as a source of nutrients for *Bacilli* (Graycar, Bott, Power, & Estell, 2013), and may face a variety of selective pressures in the extracellular environment beyond avoiding proteolytic degradation by competing proteases.

These two subfamilies (ISP and ESP) present an ideal system for investigating the energy landscapes of two proteins with the same fold, but vastly different kinetic barriers. Here, we compare the folding landscapes and dynamics of an ISP and ESP from *Bacillus subtilis*, Intracellular Subtilisin Protease 1 (ISP1) and Subtilisin E (SbtE) (Fig. 1), found in to be in the same phylogenetic clade (Subbian et al. 2004). We use several biophysical approaches to monitor both the global stability and the kinetics of folding/unfolding as well as hydrogen-deuterium exchange monitored by mass spectrometry (HDX-MS) to monitor local stability and dynamics. We find that despite their high sequence similarity and structural homology, ISP1 and SbtE have very different conformational landscapes. In addition to differences in the native state energetics and dynamics, the effects of their pro-domains are also very different. While the SbtE pro-domain both speeds up folding to the native state and provides added stability to the folded state (Kobayashi & Inouye, 1992; Li, Hu, Jordan, & Inouye, 1995; Zhu, Ohta, Jordan, & Inouye, 1989), the ISP1 pro-domain has minimal effects on both folding and stability. Interestingly, the differences in their global energetics are reflected in local changes within the core – particularly the central alpha helix – pointing to a potential basis for these differences. Additionally, while the ESP has a very large barrier to unfolding, in the absence of the pro-domain it is not completely rigid, showing a greater degree of hydrogen exchange in the core than anticipated – contrary to findings for other kinetically-trapped, secreted proteases.

**Figure 1:**
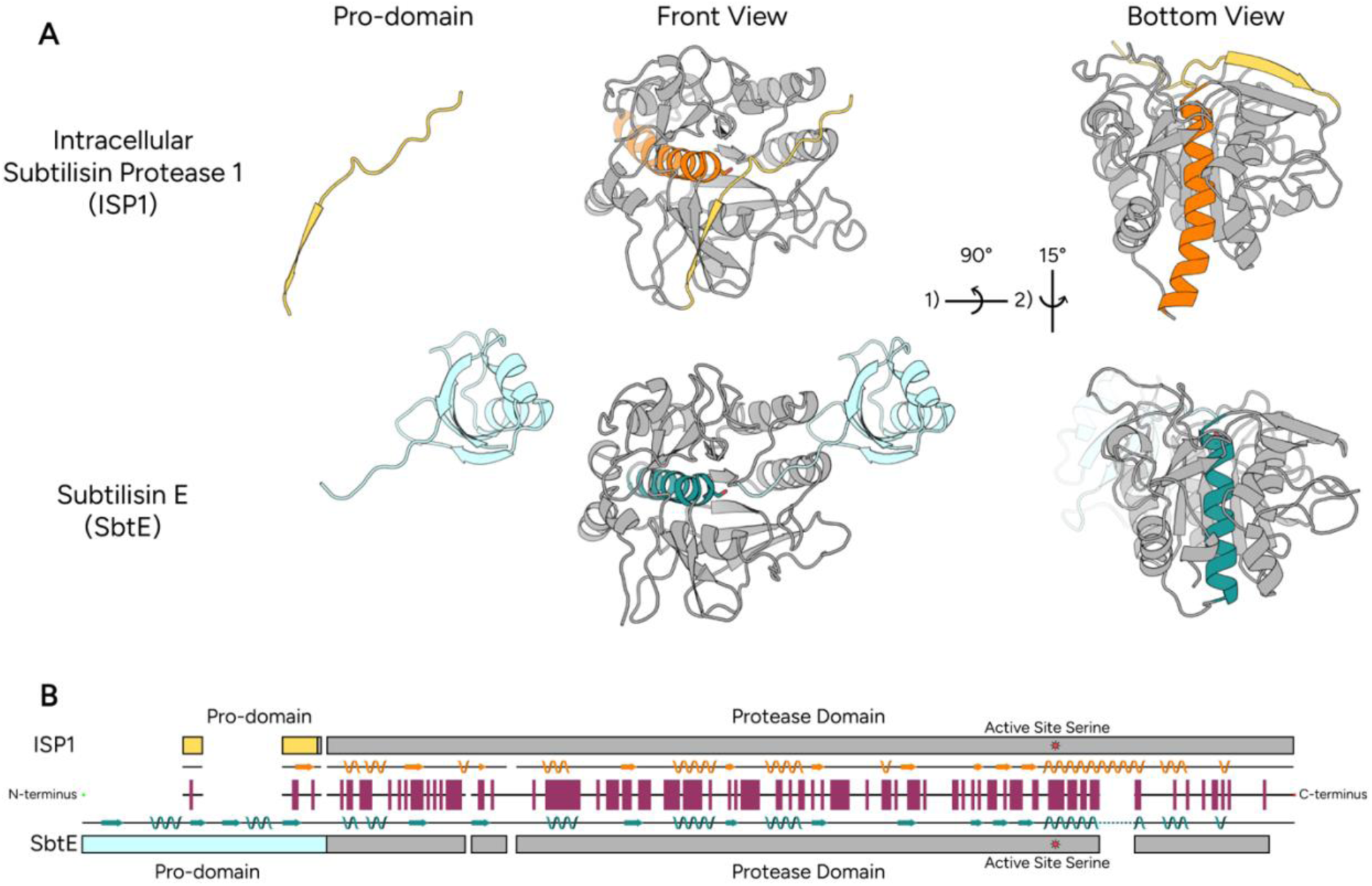
Comparison of the structures and sequences of intracellular and extracellular subtiliases. **A)** Ribbon representation of ISP1, with the pro-domain shown in yellow the core alpha helix shown in orange, and the active serine side chain as a stick, seen from the front and bottom. Residues 301-319 are not shown. The pro-domain alone is shown to the left. The structure was generated with AlphaFold3 (Abramson et al., 2024). Ribbon representation of SbtE, with the pro-domain shown in light blue, the core alpha helix shown in teal, and the active site serine side chain as a stick, seen from the front and bottom (PDB entry 1SCJ). The pro-domain alone is shown to the left. **B)** Representation of sequence alignment of ISP1 and SbtE – the pre-sequence from SbtE was not included in the alignment. The length of the alignment is represented by the black line from left to right with the N-terminus shown in green and C-terminus shown in red. Breaks in the line represent insertions into either ISP1 (first three breaks) or SbtE (remaining breaks). Beta sheets and alpha helices present in each protein are represented above and below the alignment. Any secondary elements spanning a break are connected by a dashed line. The ISP1 pro-domain is marked by the light orange box at the top and the SbtE pro-domain is marked by the light blue box at the bottom. ISP1 and SbtE have 45.7% sequence identity and conserved residues are denoted by orange boxes along the sequence.

## Results

### Intracellular Subtilisin Protease 1 (ISP1) Conformational Landscape: Global stability and re/unfolding kinetics

We first evaluated the global stability and folding/unfolding kinetics of the Intracellular Subtilisin Protease 1 (ISP1), both the mature and pro-protein. To avoid auto-proteolysis, we generated inactive variants by changing the active-site serine (residue 250) to alanine in both variants (ISP1 S250A and Pro-ISP1 S250A). Global stability was determined by carrying out equilibrium chemically induced denaturation monitored by following the circular dichroism (CD) signal at 222 nm as a function of guanidinium chloride (GdmCl) for both Pro-ISP1 S250A and ISP1 S250A. For both proteins, denaturation was reversible; the results for each denaturant point were independent of the history of the sample.

The denaturation profile of ISP1 S250A showed two cooperative transitions, one with a midpoint near 0.93 ± 0.02 M GdmCl and the other with a midpoint near 1.92 ± 0.02 M GdmCl (Fig. 2A). These data were fit to a three-state model (U ⇋ I ⇋ N) with linear extrapolation (Barrick & Baldwin, 1993) resulting in a ΔG_NI_ = 5.88 ± 0.36 kcal•mol^−1^ and ΔG_IU_ of 1.86 ± 0.50 kcal•mol^−1^, summing to a ΔG_NU_ of 7.74 ± 0.86 kcal•mol^−1^ (Table 1). The denaturation profile of Pro-ISP1 S250A also revealed two transitions, one with a midpoint near 1.1 ± 0.08 M GdmCl and the other with a midpoint near 1.7 ± 0.33 M GdmCl (Fig. 2A). The data were again fit to a three-state model (ΔG_NI_ = 7.34 ± 0.29 kcal•mol^−1^, ΔG_IU_ of 1.23 ± 0.21 kcal•mol^−1^, summing to a ΔG_NU_ of 8.57 ± 0.21 kcal•mol^−1^) (Table 1). Thus, the presence of the pro-domain has minimal effect on the global energetics of ISP1.

**Figure 2:**
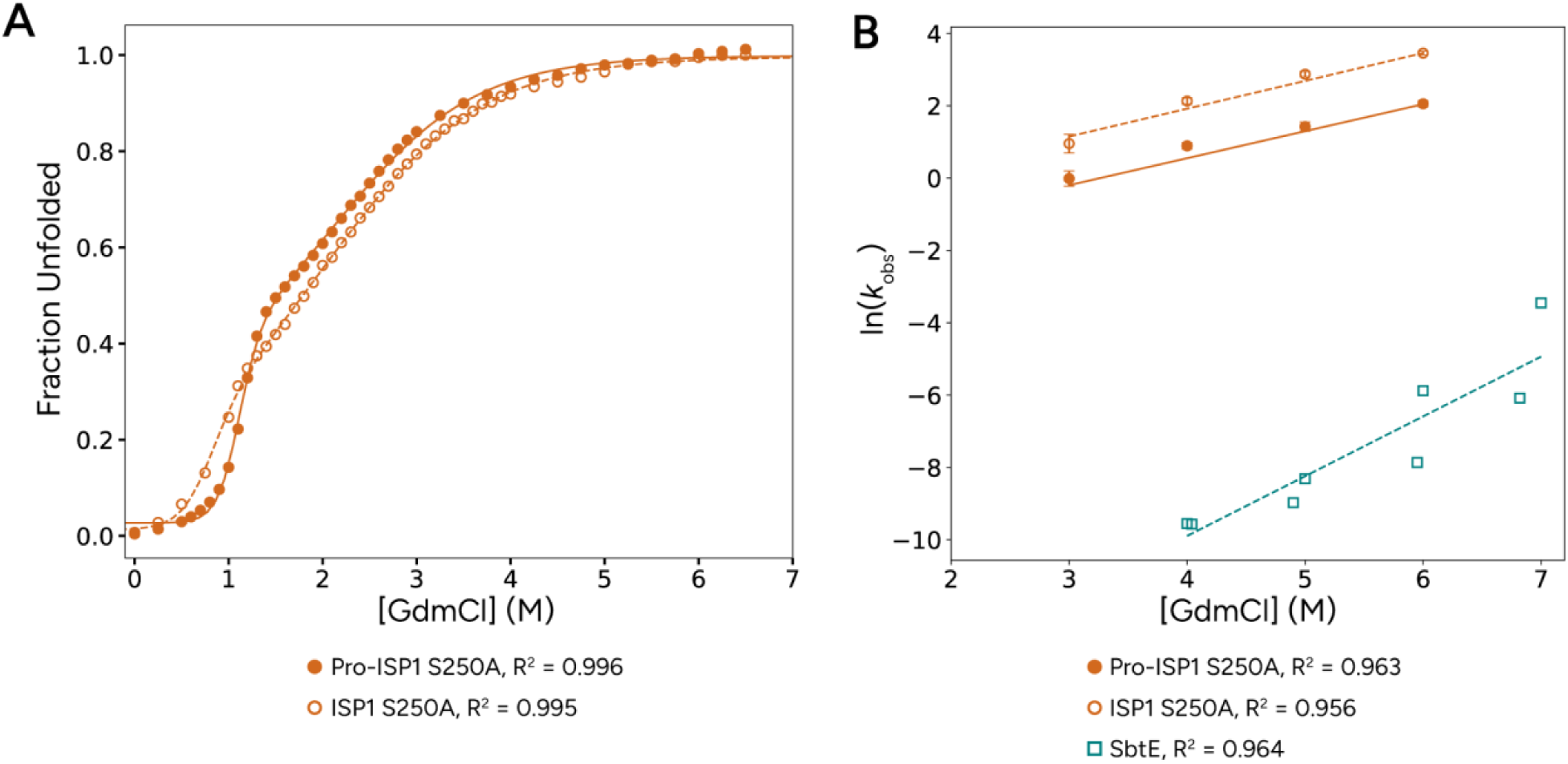
Equilibrium denaturation and unfolding kinetics demonstrate that ISP1 behaves as a typical thermodynamically stable protein and SbtE is slow to unfold. **A)** Representative guanidinium denaturation curves of Pro-ISP1 S250A (filled circles) and ISP1 S250A (open circles) monitored by CD at 222 nm. Three-state fits for Pro-ISP1 S250A (solid line) and ISP1 S250A (dashed line) are shown. **B)** Unfolding as a function of guanidinium concentration for Pro-ISP1 S250A (filled circles, orange), ISP1 S250A (open circles, orange), and SbtE in the presence of PMSF (open squares, blue) as monitored by the CD signal at 222 nm. Linear regressions for Pro-ISP1 S250A (solid line, orange), ISP1 S250A (dashed line, orange), and SbtE (dashed line, blue) are shown. All data represented here was collected at pH 8 (see *Materials and Methods)*.

**Table 1:** Thermodynamic and kinetic properties of ISP1 S250A, Pro-ISP1 S250A, SbtE, and Pro-SbtE.

|  | *ISP1 S250A* | *ISP1 S246A<sup>a</sup>* | *Pro-ISP1 S250A* | *SbtE* | *Pro-SbtE S221C<sup>a</sup>* |
| --- | --- | --- | --- | --- | --- |
| $\Delta G_{NI} (kcal\ mol^{-1})$ | $5.88 \pm 0.36$ | 3.56 | $7.33 \pm 0.05$ | NA | 1.62 |
| $m_{NI} (kcal\ mol^{-1}\ M^{-1})$ | $6.36 \pm 0.51$ | 7.14 | $6.93 \pm 0.50$ | NA | 1.11 |
| $\Delta G_{IU} (kcal\ mol^{-1})$ | $1.86 \pm 0.50$ | 1.74 | $1.23 \pm 0.29$ | NA | 8.36 |
| $m_{IU} (kcal\ mol^{-1}\ M^{-1})$ | $0.93 \pm 0.02$ | | $0.735 \pm 0.027$ | NA | |
| $\Delta G_{NU} (kcal\ mol^{-1})$ | $7.74 \pm 0.86$ | 5.3 | $8.56 \pm 0.34$ | <-3.73** | 9.98 |
| $m_{NU} (kcal\ mol^{-1}\ M^{-1})$ | $4.2 \pm 0.5$ | 7.85 | $6.1 \pm 0.2$ | | 3.02 |
| $k_U (s^{-1})$ | $1.43E-1 \pm 5.21E-2$ | 5E-2<sup>b</sup> | $8.85E-2 \pm 7.33E-3$ | $1.92E-7 \pm 1.27E-7^{pH\ 8}$,<br>$2.8E-5 \pm 1.9E-5^{pH\ 5}$ | 1E-1<sup>b</sup> |
| $k_f (s^{-1})$ | >180 | 20.23<sup>c</sup>,<br>0.012<sup>d</sup> | >180 | <4.92E-8*<sup>25°C</sup>,<br><5.07E-8*<sup>4°C</sup> | 20.23, 1.2E-2 |

\*Estimated base on lower limit of detection for refolding by activity, \*\*Calculated based on the ratio of the kinetic measurements, (a) Estimated based on the Chevron previously reported (Subbian et al., 2004), ISP1 S250A uses the crystal structure numbering and ISP1 S246A uses the sequence numbering, (b) Estimate based on previously reported chevron (Subbian et al., 2004), (c) Previously reported $k_{UI}$ (Subbian et al., 2004), and (d) Previously reported $k_{IN}$ (Subbian et al., 2004). All experiments were carried out at 25°C in 50 mM Tris pH 8, 500 mM (NH<sub>4</sub>)<sub>2</sub>SO<sub>4</sub>, 1 mM CaCl<sub>2</sub>, 0.5 mM TCEP, unless otherwise specified, and reported values are the average of three to four independent repeats. SbtE refolding was
carried out in 50 mM Bis-Tris pH 5, while unfolding was carried out in 50 mM Potassium Acetate pH 5, both with 500 mM $(\text{NH}_4)_2\text{SO}_4$ , 1 mM $\text{CaCl}_2$ , 0.5 mM TCEP.

To assess ISP1’s kinetic barrier in the absence and presence of the pro-domain we monitored the time course of refolding and unfolding for the same variants initiated by either diluting into (unfolding) or out of (refolding) high concentrations of guanidinium (see *Materials and Methods)*. For both ISP1 S250A and Pro-ISP1 S250A, the entire refolding process takes place within the deadtime of the instrument (5.6 milliseconds for stopped-flow fluorescence and ∼15 seconds for manual mixing and CD). Thus, the rate of refolding is greater than 180 s^−1^ (τ_unfolded_ of less than 5.6 ms). In contrast, unfolding of ISP1 S250A and Pro-ISP1 S250A shows observable kinetics when monitored by fluorescence that fit to a single exponential accounting for the entire signal change (Fig. S1). Linear extrapolation of ln*k*_u_ as a function of GdmCl concentration (Fig. 2B) results in an unfolding rate of 0.143 ± 0.0521 s^−1^ for ISP1 S250A and 0.0885 ± 0.00898 s^−1^ for Pro-ISP1 S250A (Table 1). Thus, the presence of the pro-domain does not appear to impact the unfolding rate for ISP1. In sum, the pro-domain has minimal effects on the folding stability and kinetic for ISP1.

### Subtilisin E (SbtE) Conformational Landscape

Next, we evaluated both the global stability and folding/unfolding kinetics of the extracellular Subtilisin E (SbtE). As expected, the refolding and unfolding of SbtE is too slow to allow determination of global stability using standard equilibrium experiments. Instead, we turned to kinetic measurements to monitor the unfolding and folding rates and used these to obtain an estimate for the global stability. To generate mature SbtE, we used wild-type (active) SbtE, which when expressed with the pro-domain, auto processes to yield mature SbtE (see *Materials and Methods*).

Unfolding of mature SbtE was monitored by CD, with addition of the inhibitor PMSF to prevent autodigestion during the unfolding process (Chu, Chao, & Bi, 1995; Ikemura & Inouye, 1988), at pH 5 or 8 (See *Materials and Methods*). The unfolding rate was measured at pH 5 for comparison with SbtE refolding experiments described below and at pH 8 for comparison with ISP1 S250A and Pro-ISP1 S250A unfolding experiments. SbtE unfolding traces were fit to a single exponential (Fig. S2 and S3) and linear extrapolation of ln*k*_u_ as a function of guanidinium concentration (Fig. 2B), which results in an estimate for *k*_u_ in the absence of denaturant of 1.92×10^−7^ ± 1.27×10^−7^ s^−1^ at pH 8 (Table 1). SbtE unfolding is at least 5 orders of magnitude slower than both ISP1 S250A and Pro-ISP1 S250A. A linear extrapolation of ln*k*_u_ as a function of guanidium concentration at pH 5 results in a *k*_u_ of 2.8×10^−5^ ± 1.9×10^−5^ s^−1^ (Table 1). Thus, when compared to the unfolding rate measured for the inactive protease in the presence of the pro-domain provide *in trans* (Subbian et al., 2004), unfolding of the mature active protease alone is slower by 6 orders of magnitude.

The refolding of mature SbtE is too slow to be monitored by traditional spectroscopy. However, activity is a much more sensitive assay for folded, native protein, and therefore the concentration of active protease can be used as a proxy for the fraction folded protease. With this approach, the refolding rate can be indirectly determined by measuring the activity over time (Baker & Agard, 1994a; Baker, Sohl, & Agard, 1992b; Sohl et al., 1998; Takagi, Maeda, Ohtsu, Tsai, & Nakamori, 1996). We diluted unfolded SbtE into folding conditions and assayed for protease activity as a function of incubation time (see *Materials and Methods*). Even using this more sensitive assay, we were unable to detect activity after six days in the absence of the pro-domain. Our assay has a lower detection limit of 2×10^−10^ M folded protein (based on a standard curve using active protease), which allows us to place an upper limit of 2×10^−10^ M folded protein for every time point. However, when we assay for the total protein, [I], over time (again using an activity assay after the addition of the pro-domain), we see a decrease in protein over time, presumably due to autodigestion over the long incubation time coming from the small (undetectable) amount of folded protein, [N] (Fig. 3, see *Materials and Methods*). Thus, we can use our upper limit of folded protein normalized by the determined concentration of total protein to calculate an upper limit of fraction folded as a function of time. These data were then analyzed using the model derived by Sohl et al. 1998 to generate an estimate of the folding rate. Given all the assumptions noted above, this rate (*k*_f_ of 4.92×10^8^ s^−1^ at 4°C and 5.07×10^8^ s^−1^ at 25°C, Table 1) represents an upper limit on the folding rate for SbtE (folding of SbtE must be slower than ∼5×10^8^ s^−1^). Comparing these rates to previously published rates for Pro-SbtE (Subbian et al., 2004), the pro-domain must increase the refolding rate by at least 9 orders of magnitude.

**Figure 3:**
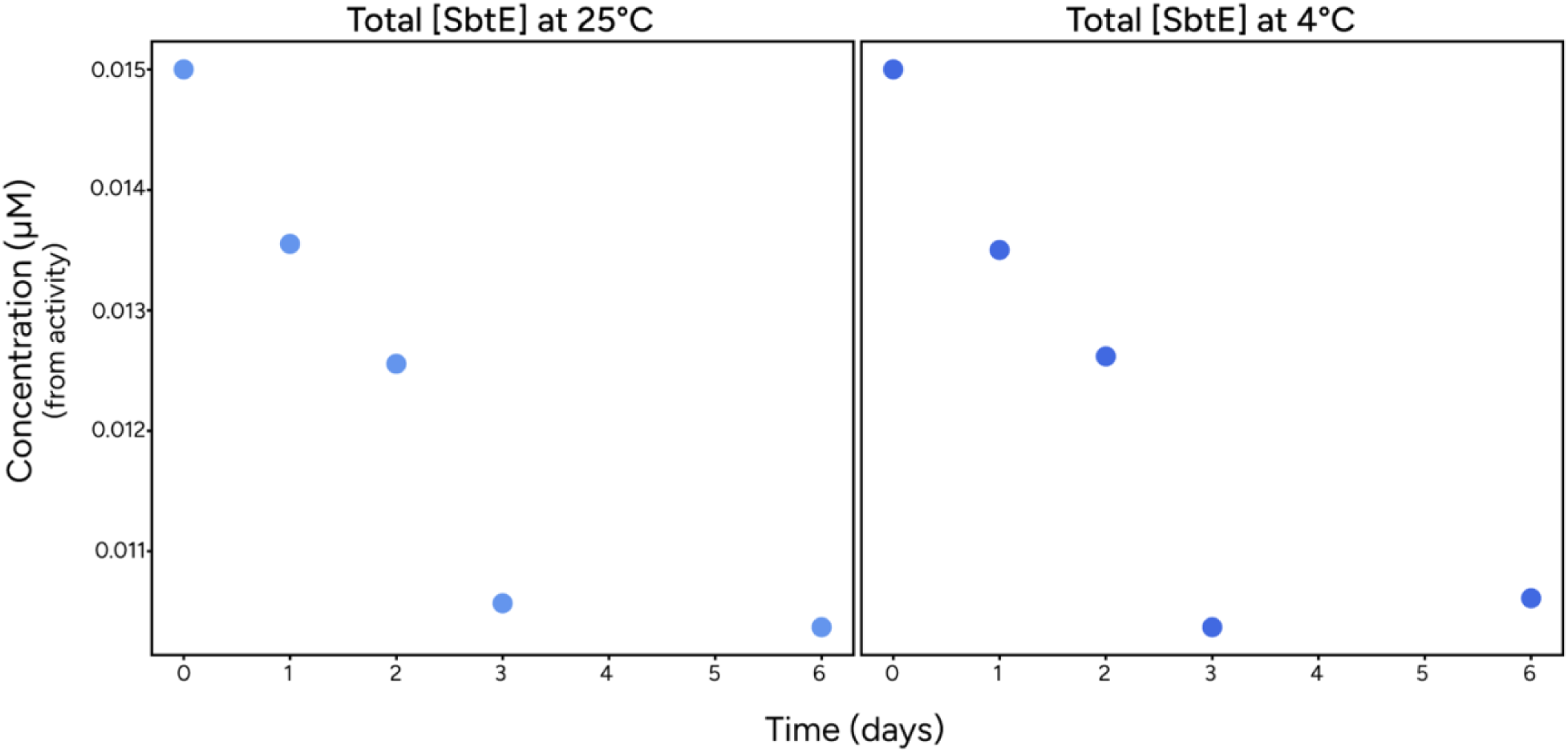
The concentration of total SbtE as a function of refolding time. Mature SbtE was diluted into refolding buffer in the presence of phenylboronic acid and the total concentration of activity was monitored as a function of time by adding an excess of pro-domain and then assayed for activity. A direct activity assay on the incubated protein was showed no detectable activity, putting an upper limit of 2×10^−10^ M folded protein at every timepoint. Together, this upper limit of activity and measured amount of total protein allowed an upper estimate of the folding rate of SbtE of 4.92×10^8^ s^−1^ at 4°C and 5.07×10^8^ s^−1^ at 25°C (see *Materials and Methods*).

### Local Stability by HDX-MS

To investigate the effects on the energy landscape at a more local level, we turned to hydrogen-deuterium exchange monitored by mass spectrometry (HDX-MS), first on the mature proteins (ISP1 S250A and SbtE) and then on the pro-proteins (Pro-ISP1 S250A and Pro-SbtE S221A). We monitored the exchange process in a continuous exchange experiment (25°C, pH 8, see *Materials and Methods*), following the uptake of deuterium as a function of time (Fig. 4A). HDX was monitored using mass spectrometry at the level of peptides from time points as short as 15 s to as long as 1 year (see *Materials and Methods*). Both proteins digest well with >95% coverage of the protein sequence and > 500 peptides from ISP1 S250A and > 150 peptides from SbtE (Fig. S4) allowing us to measure HDX across the whole protein.

**Figure 4:**
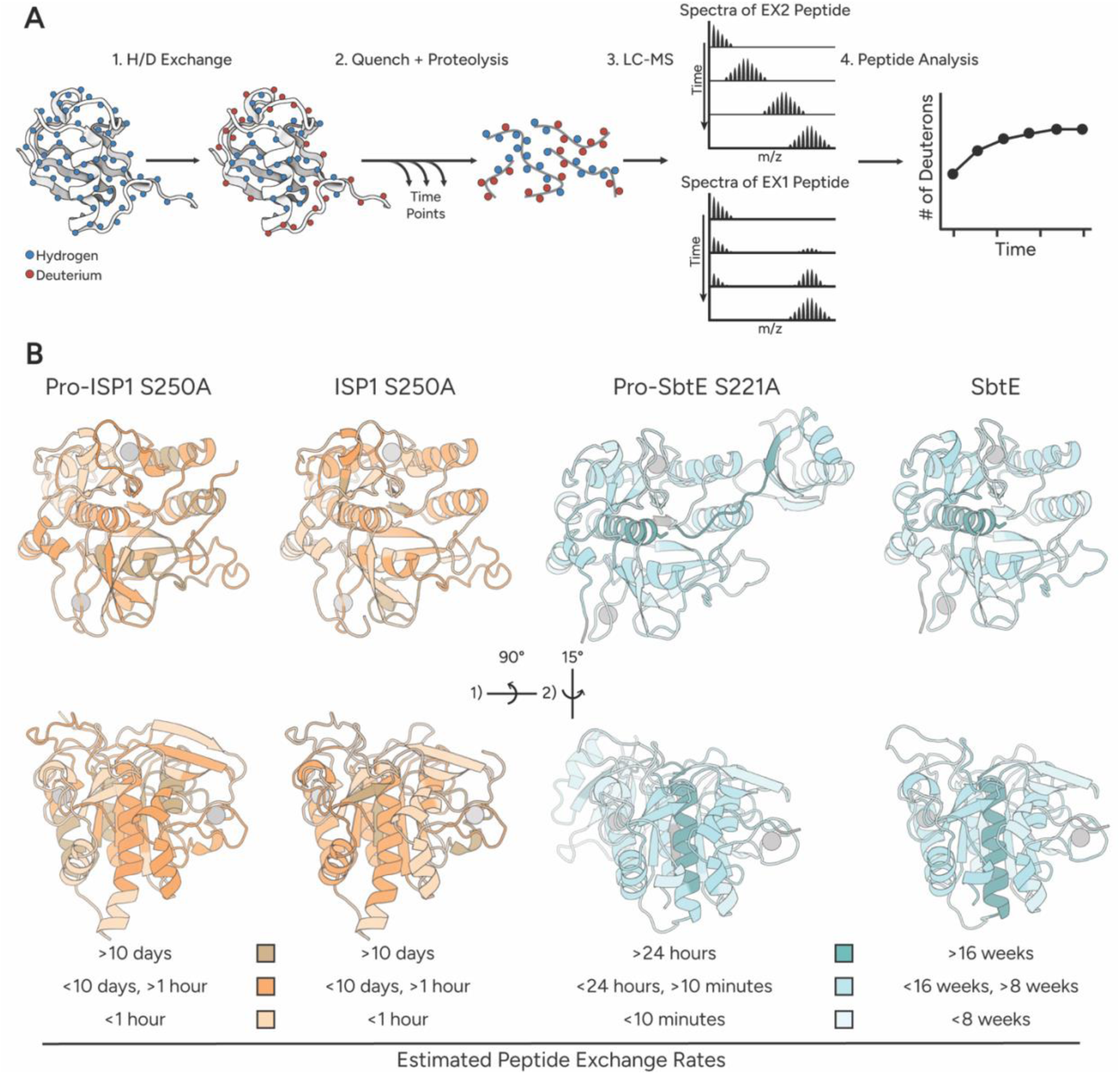
HDX-MS measures local protein dynamics. **A)** HDX-MS workflow. Protein is incubated in deuterated buffer and aliquots are quenched over time. Quenched samples are subjected to proteolysis and run on an LC-MS. EX2 behavior is characterized by a shifting mass spectrum, while EX1 behavior shows a bimodal mass spectrum. Mass spectra are analyzed and the number of deuterons taken up by a peptide is plotted as a function of time. **B)** Ribbon representations of Pro-ISP1 and ISP1 in orange and Pro-SbtE and SbtE in teal are seen from the front and bottom. For simplicity, only peptides less than 15 amino acids in length are shown. Colors reflect their estimated hydrogen exchange rate as determined by the half-time for each peptide (time at which half the amide protons exchanged). Calcium ions are shown as gray circles.

The data were analyzed using the standard Linderstrøm-Lang model (Wildes & Marqusee, 2004), where hydrogen exchange occurs under one of two kinetic regimes, so-called EX1 and EX2, that can be easily differentiated by mass spectrometry (Fig. 4A). Each report on a different aspect of the fluctuations that allow exchange. In EX1, the observed rate of exchange reports on the kinetics of this fluctuation or opening event (slower exchange indicates a larger kinetic barrier to the exchange competent state) and in EX2 the observed rate of exchange reports on the fraction of the population in the exchange-competent or open state (thermodynamics, slower exchange implies a rarer population (higher energy) for the exchange competent state). All exchange reactions were conducted at pH 8. When monitored by mass spectrometry, EX2 exchange results in a single peak that shifts from a lighter to heavier average m/z over time, while EX1 results in two peaks (bimodal behavior) where the intensity of the lighter peak decreases over time as the intensity of heavier peak increases (Fig. 4A).

The overall pattern of hydrogen exchange is similar for all four proteins (Fig. 4B) consistent with their overall fold. In general, the cores of the protein are the most protected with the loops less protected. The measured rates of exchange, however, are quite different. For instance, the two mature proteins show very different hydrogen exchange rates (Figs. Fig 4B, and Fig. 5A). On the whole, ISP1 S250A displays HDX behavior expected for a thermodynamically stable protein. Most of the peptides show EX2 exchange that is complete within two weeks, consistent with the expected global stability of 7 kcal/mol. SbtE, however, showed much more complicated HDX behavior. While the majority of peptides examined do show EX2 behavior – we observed only minimal exchange over the time course of the experiment. Thus, the observed rate of exchange was very slow (not reaching full exchange even after one year), too slow to arise from the unfolded state based on the stability estimates in Table 1 and suggesting that these exchange events arise from very high energy (rare) fluctuations on the native side of the barrier.

**Figure 5:**
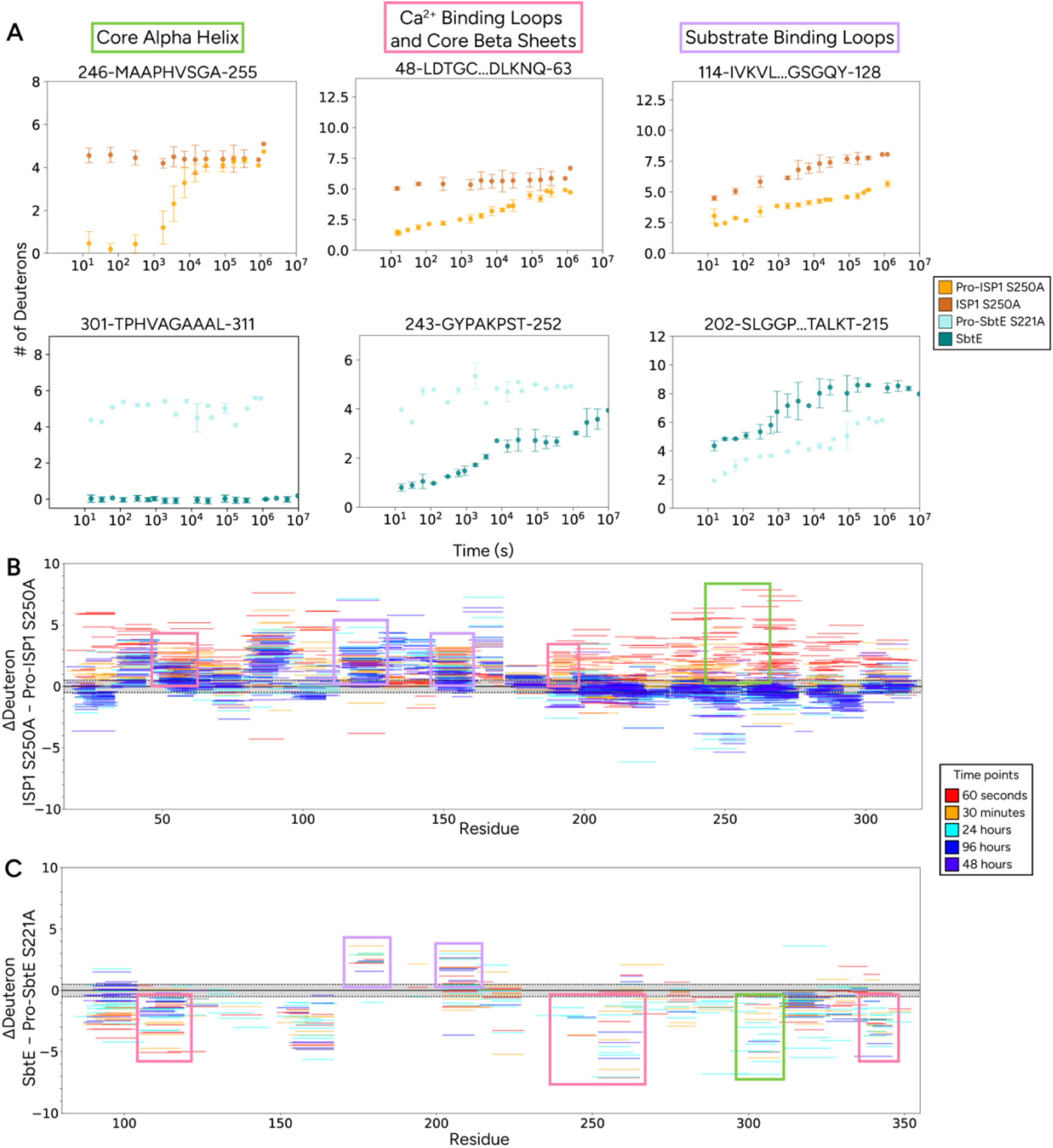
HXD-MS analysis shows local dynamics are modulated by the pro-domain in opposite manners for ISP1 and SbtE. **A)** Uptake plots showing average number of deuterons per protein as a function of time for representative peptides (average of three replicates, error bars represent standard deviation not back exchange corrected). Plots are colored coded by protein: Pro-ISP1 S250A in orange, ISP1 S250A in dark orange, Pro-SbtE S221A in light blue, SbtE in dark blue. Pro-ISP1 S250A’s core helix exchanges with a *k*_obs_ of ∼10^−4^ s^−1^, ISP1 S250A and Pro-SbtE S221A’s core helix exchanges with a *k*_obs_ of >10^−1^ s^−1^. ISP1 S250A’s core helix exchanges with a *k*_obs_ of >10^−1^ s^−1^ and SbtE’s core helix exchanges with a *k*_obs_ of <10^−8^ s^−1^. Pro-ISP1 S250A’s calcium binding loops and core beta sheets exchange with a *k*_obs_ of ∼10^−5^-10^−6^ s^−1^. ISP1 S250A’s calcium binding loops and core beta sheets exchange with a *k*_obs_ of >10^−1^ s^−1^. Pro-SbtE’s calcium binding loops and core beta sheets all exchange with a *k*_obs_ of >10^−1^ s^−1^. SbtE’s calcium binding loops and core beta sheets exchange with a *k*_obs_ of ∼10^−5^-10^−7^ s^−1^. Pro-ISP1 S250A’s substrate binding loops exchange with a *k*_obs_ of ∼10^−1^ s^−1^ and ISP1 S250A’s exchange with a *k*_obs_ of ∼10^−5^ s^−1^. Pro-SbtE S221A’s substrate binding loops exchange with a *k*_obs_ of ∼10^−4^ s^−1^ and SbtE’s exchange with a *k*_obs_ of ∼10^−1^ s^−1^. Additonal uptake plots for pepitdes with the same behavior can be found in Fig. S5). **B)** Difference Woods plots for ISP1 S250A vs Pro-ISP1 S250A and **C)** SbtE vs Pro-SbtE S221A. Each peptide is plotted, and color coded by time point: 60 seconds in red, 30 minutes in orange, 24 hours in cyan, 48 hours in blue, and 96 hours in purple. Positive values indicate more deuteration (less protection) in the pro-proteins and negative values indicate less deuteration (more protection) in the first protein listed. Regions corresponding to uptakes plots in b are boxed.

Analysis of the individual peptides allowed us to interpret these fluctuations at a more local level, where there are interesting differences in behavior, most notably in the core alpha helix of the proteins (Fig. 1A). For ISP1 S250A, peptides in this helix exchange rapidly (in less than 15 s), despite arising from the central core of this thermodynamically stable protein (Fig. 5A). Thus, this core region is surprisingly flexible, and exchange is not limited by global unfolding of the protein. In contrast, in SbtE, peptides from this region show no observable exchange, even out to a year (Fig. 5A, Fig. S8), consistent with a model where exchange from this region is likely in EX1 and requires crossing the high-energy kinetic barrier to global unfolding. The core beta sheets and calcium-binding loops of SbtE, however, show very slow EX2 exchange (Fig. 5A, Fig. S9), consistent with a rigid core of the protein that still undergoes some fluctuations to exchangeable conformations. In sum, while ISP1 S250A’s exchange pattern suggests a high degree of flexibility in the core-region of the protein, SbtE’s core exchange pattern suggests a high degree of rigidity.

Next, we carried out HDX on the proteins containing the pro-domain using the inactive variants Pro-ISP1 S250A and Pro-SbtE S221A. We monitored continuous HDX at 25°C, pH 8 from time points as short as 15s to as long as 2 weeks. Again, both proteins are amenable to HDX-MS with >95% coverage of the protein sequence and >700 peptides from Pro-ISP1 S250A and >400 peptides from Pro-SbtE S221A (Fig. 4B, Fig. S4). Both pro-proteins behaved like typical thermodynamically stable proteins, exhibiting EX2 exchange throughout the entire structure with exchange rates consistent with thermodynamically stable proteins. For ISP1, compared to the mature protein, the addition of the pro-domain slows hydrogen exchange (Fig. 5B), consistent with the overall increase in global stability (Table 1). There are also notable differences at the local level. In particular, Pro-ISP1 S250A’s core alpha helix (Fig. 1A) shows slowed exchange consistent with exchange from the globally unfolded state. In sum, these data suggest that compared to the mature ISP1 S250A, the pro-protein behaves more like a typical folded protein with a rigid core.

In contrast, for Pro-SbtE S221A the presence of the pro-domain increases the observed hydrogen exchange compared to SbtE (Fig. 5B). The one exception to this observation is at the active-site loops that contact the pro-domain (Fig. 5A, Fig. S6), as would be expected from the increased burial of these residues. The core alpha helix, which in the absence of the pro-domain appears to exchange from the globally unfolded state, shows increased exchange and therefore increased flexibility when the pro-domain is present (Fig. 5A). Thus, unlike the intracellular protein, the presence of the pro-domain increases the fluctuations from this core alpha helix. In sum, removal of the pro-domain results in a decrease in the overall flexibility of SbtE, which is consistent with conversion of a thermodynamically stable protein to a kinetically trapped protein (Jaswal, Sohl, Davis, & Agard, 2002b).

### Increased local stability and proteolytic resistance

As mentioned above, core beta sheets in SbtE show slow exchange – this exchange is in the EX2 kinetic regime as shown by a shifting centroid (Fig. S9) and under thermodynamic control. On the other hand, the core alpha helix may be in the EX1 kinetic regime and under kinetic control. The presence of equilibrium fluctuations in SbtE differs from αLP, which was shown to only be under kinetic control (Jaswal et al., 2002a).

To determine if the decreased fluctuations seen in the core of the mature protein, SbtE, correlate with increased proteolytic resistance, as with αLP (Jaswal et al., 2002a), we carried out proteolytic competition assays using thermolysin as a model of a thermodynamically stable protease. The two proteases (SbtE and thermolysin) were added in equimolar amounts and the amount of each protease remaining was monitored as a function of time over a 24 hour time period. Proteolysis was monitored by SDS-PAGE gel analysis (see *Materials and Methods*). Thermolysin and subtilisins have similar substrate preferences, both showing a preference for bulky, hydrophobic amino acids (Heinrikson, 1977; Keil, 1992; Takagi et al., 1996; J A Wells, Cunningham, Graycar, & Estell, 1987). Figure 6 shows that digestion of thermolysin is nearly completely within 15 s while SbtE remains undigested out to 24 hrs, >10,000 times longer. Thus, like αLP, SbtE’s kinetic stability provides proteolytic resistance as compared to thermolysin. Therefore, the observed fluctuations in the core are not sufficient for degradation. As expected, based on its thermodynamic stability (therefore low population of unfolded protein), ISP1 S250A is resistant to degradation in this time frame (data not shown). In contrast to the competition with thermolysin, SbtE and ISP1 have similar proteolytic resistance, one through kinetic control and one through thermodynamic control.

**Figure 6:**
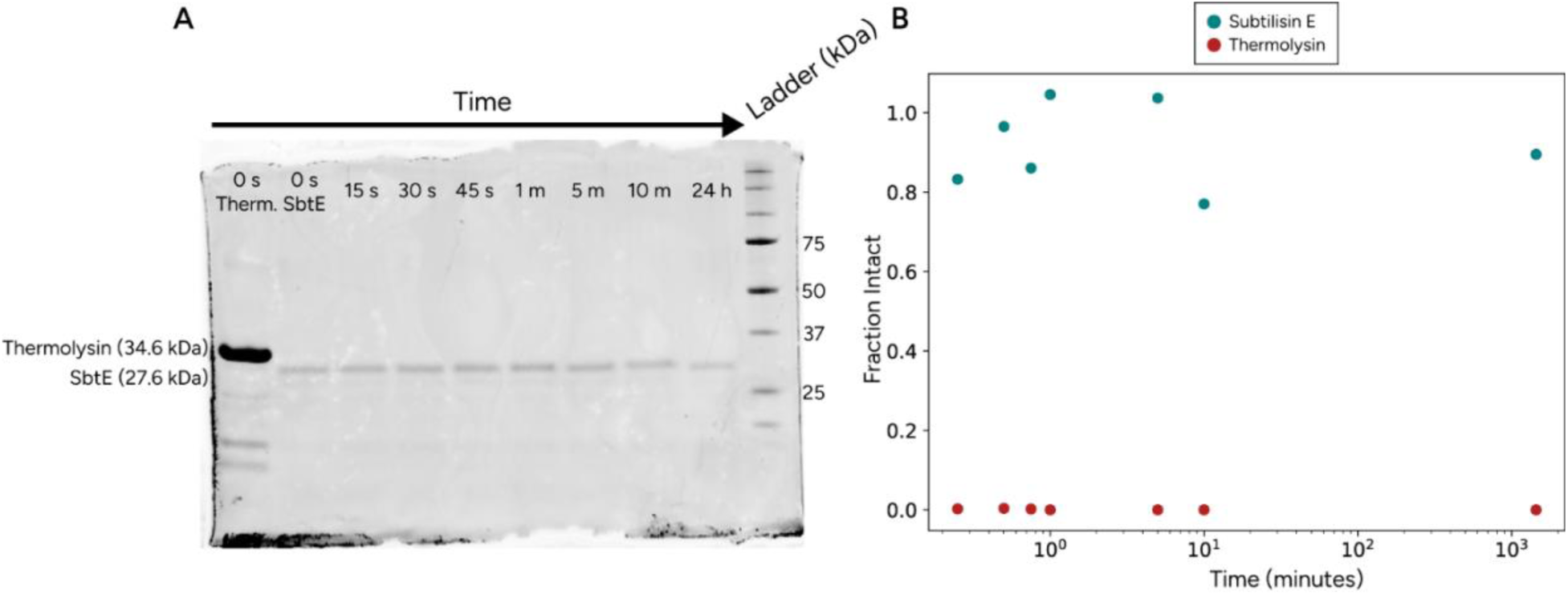
Subtilisin outlasts thermolysin in an *in vitro* competition. **A)** Representative SDS-PAGE gels showing thermolysin is rapidly degraded but SbtE persists over a 24-hour time course. **B)** Quantification of gel bands from part A. The band from each time point was normalized to their respective 0 second time point for each protease and plotted as a function of time.

## Discussion

Ever since the classic work of Anfinsen and Levinthal (Anfinsen, 1973; Levinthal, 1968), how proteins manage to navigate their vast conformational landscape to fold in a biologically relevant timescale has been an important and heavily investigated area. It is clear, however, that once folded, the lifetime of the native conformation (both its fluctuations and global unfolding) is also under selective pressure. The mechanisms that control these partial and global unfolding events and the lifetime of the folded state are not well understood. Here, we have examined two subtiliases from the same organism, with the same basic fold, that function in very different environments (ISP1 and SbtE from *Bacillus subtilis*). Our studies are based on previous analyses of ISP1 and Pro-SbtE S221C, where we now estimate and compare their energy landscapes with and without the pro-domain. From a comparison of their global energetics and dynamics as well as their local dynamics, we find that, despite sharing the same fold and a high level of sequence identity, ISP1 and SbtE have very different energy landscapes at all levels: their global stability (thermodynamics), unfolding/refolding kinetics, and fluctuations at the subglobal level are all different. While each has a pro-domain, the small pro-domain of the intracellular protein (ISP1) has very little effect on the energetics and folding of the protein. For the extracellular protein, the pro-domain serves as an apparent intramolecular chaperone, speeding up folding, and generating a thermodynamically unstable, kinetically trapped mature protein with a high barrier for unfolding. The mature SbtE shows high energy, but measurable, fluctuations from the native state, unlike the well-characterized kinetically trapped protein Alpha-lytic protease (Jaswal et al., 2002a), which is relatively rigid.

As mentioned before, other secreted proteases, like αLP, are known to be thermodynamically unstable and kinetically trapped (so called ‘kinetic stability’) and dependent on their pro-domain for folding. The biophysical studies of αLP laid the foundation for our understanding of what the energy landscape of a kinetically trapped protein may look like and why they may have evolved (Baker & Agard, 1994a; Baker et al., 1992b; Jaswal et al., 2002a; Sohl et al., 1998). In these studies, the sequence of αLP was compared with other chymotrypsin-like proteases, but as αLP has no known direct homologs, it was not subjected to the comparative analysis carried out here. Subsequent studies examining the relationship between pro-domain size and kinetic stability for extant, but inactive, αLP relatives and their ancestral states found that pro-domain size correlated with the height of the kinetic unfolding barrier and that the proteins cluster phylogenetically based on pro-domain size, providing additional insight into structural features that impact the energetics of different αLP family members (Nixon et al., 2021). The work presented here on two subtiliases is, to our knowledge, the first direct comparison of two homologs, from the same species, and the difference in their global and local energetics in the presence and absence of their pro-domains.

### Differences in global energetics and kinetics

The intracellular protease ISP1 behaves, as expected, as a typical thermodynamically stable, three-state folder, both in the presence and absence of its pro-domain. While ISP1 S250A’s folding thermodynamics and kinetics were previously characterized in Subbian et al. 2004, the impact of the pro-domain on the energetic landscape was not assessed. Like most intracellular subtiliases, ISP1 has a small (17 aa) pro-domain. Despite the evolutionary conservation of the pro-domain, we find that the pro domain has little impact on the energy landscape of ISP1. The presence of the pro domain increases the global stability of the protein by less than one kcal/mol (7.74 ± 0.86 to 8.56 ± 0.34). Similarly, the kinetic barrier for unfolding is quite low, both in the presence and absence of the pro domain (*k*_u_ = 1.43×10^−1^ ± 5.21×10^−2^ s^−1^ and 8.85×10^−2^ ± 7.33×10^−3^ s^−1^, respectively). Thus, for the intracellular protease, the pro-domain (only 17 amino acids) does not appear to function as an intramolecular chaperone, but more likely as a regulator for the proteolytic activity of the enzyme (zymogen) – which supports the previously untested hypothesis that the pro-domain plays a regulatory role.

In order to determine the role of the pro-domain, we replicated the equilibrium and kinetic experiments for ISP1 S250A in our buffer containing reducing agent, TCEP, and found slight differences in the resulting global stability and refolding kinetics (Subbian et al. 2004). As an intracellular protein, ISP1 normally exists in a reducing environment, ensuring that the two cysteines present in the protein remain in a reduced oxidation state. We find that ΔG_NI_ is ∼2.32 kcal/mol higher than the previously reported value measured in the absence of reducing agent, with no difference in ΔG_IU_. Additionally, we see an ∼10 fold difference in the refolding rate, with no notable difference in the unfolding rate (Table 1) (Subbian et al., 2004). These small changes may suggest that the unfolded state is stabilized in an oxidizing environment, but that the intermediate and native state are unaffected. The two cysteines in ISP1 are located distant to each other in the native state and are unlikely to form a disulfide bond.

The pro-domain plays a very different role in the extracellular protease. For SbtE, the pro-domain has a large impact on the energy landscape of the protease - this 78 amino acid pro-domain increases the stability of the protein. Although we cannot precisely determine the global stability of mature SbtE (because we are unable to detect any folded protein (by activity) within our experimental limits), we can conclude that the protein is not thermodynamically stable because the unfolding rate is faster than the folding rate (Fig. 7). It is formally possible that the inhibitors present in the refolding and unfolding reactions may stabilize the native state, which would mean that in the absence of inhibitor unfolding would occur even faster, which would only strengthen our conclusion that the protein is thermodynamically unstable. Based on previously reported data on Pro-SbtE S221C, we can estimate the pro-domain increases the stability of the protein by at least 13 kcal/mol (from an estimated maximum of 3.73 kcal/mol to almost −10 kcal/mol) (Subbian et al., 2004). Thus, like αLP, SbtE is kinetically trapped.

**Figure 7:**
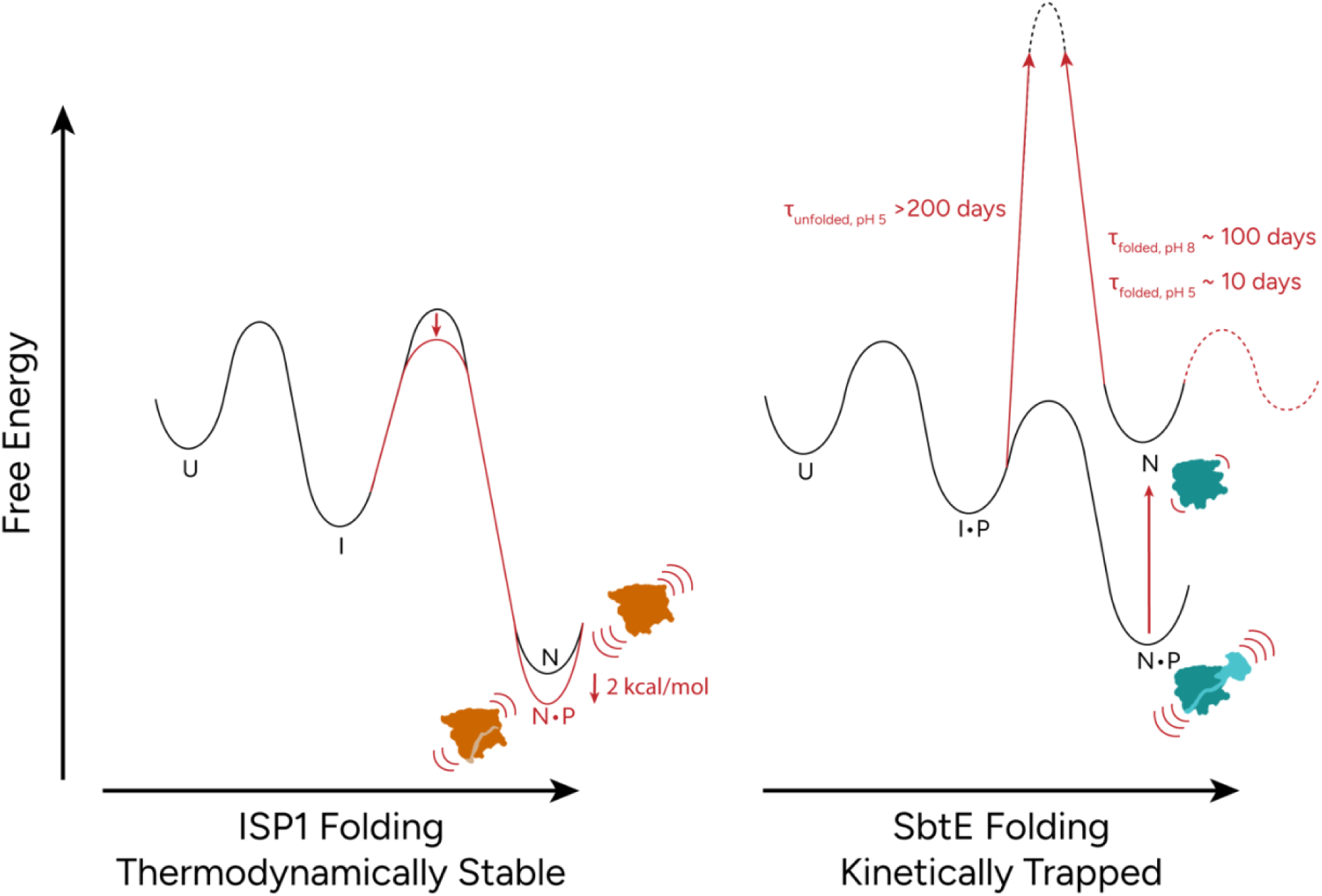
Cartoon of the energy landscapes for SbtE and ISP1. Left: ISP1 illustrating the thermodynamic stabilizing effect of the pro-domain on the native state (red lines) and decrease in local fluctuation in the presence of the pro-domain (orange). Right: SbtE illustrating the thermodynamic stabilizing effect of the pro-domain and kinetic trapping in the absence of the pro-domain (red lines). A new state, in equilibrium with the native state, is shown on the folded side of the barrier. The increase in fluctuations in the presence of the pro-domain is shown in the blue cartoon structures.

Although kinetically trapped, SbtE’s unfolding barrier does not appear to be as dramatically high as seen for αLP – with an unfolding rate not that different from many thermodynamically stable proteins (a folded state lifetime of 100 days vs 1.2 years for αLP). However, the lifetimes of both proteins are likely well beyond the biologically relevant timescale – questioning the evolutionary pressures or need for such unusually high barriers.

### Differences in local fluctuations

In addition to differences in their global energetics, the mature SbtE and ISP1 also show very different fluctuations at the local level. We found that ISP1 behaves like a typical thermodynamically stable protein with fluctuations from the native state observable by hydrogen exchange. These fluctuations are all in the EX2 regime – showing a protein with fluctuations that span in energy from near the native state to the globally unfolded protein. HDX rates are slowed in the presence of the pro-domain, rigidifying the protein as might be expected by the binding of a small domain or region (Fig. 7). Since the pro-domain is widely believed to act solely as an inhibitor to ISP1’s proteolytic activity, a reduction in fluctuations (which may facilitate activity) due to the pro-domain is reasonable.

The pro-domain has very different effects on the local dynamics of the extracellular protein, SbtE. When monitored by HDX, mature SbtE shows minimal fluctuations, whereas the Pro-SbtE S221A behaves like a typical folded protein, with fluctuations that span in energy from near the native state (ΔG_HDX_ ≤0.1 kcal/mol) to the globally unfolded protein (ΔG_HDX_ ∼6 kcal/mol). While previously reported data for Pro-SbtE thermodynamics and kinetics were measured on Pro-SbtE S221C, which mimics the cleaved complex, our HDX was carried out on Pro-SbtE S221A, which mimics the cis-folded complex, where the covalent attachment between the pro- and protease domains is still present. This difference in complex may explain why no peptides were observed to have a ΔG_HDX_ ∼10 kcal/mol, corresponding to the ΔG_NU_ of 9.98 kcal/mol reported for Pro-SbtE S221C.

While the increased fluctuations in the presence of SbtE’s pro-domain was expected (the pro-domain lowers the barrier between states making excursions from the native state more accessible), we were surprised by the extremely slow hydrogen exchange we do observe in SbtE. Based on the *k*_int_ for the peptides in this region the ΔG of HDX for the amides in these peptides would be approximately 6.5 kcal/mol, which is greater than the global stability of the protein (ΔG_NU_). This suggests that the folded state is in equilibrium with a state other than the unfolded state (Fig. 7). Given the high kinetic barrier to unfolding, these equilibrium fluctuations must arise from the native side of the unfolding barrier.

The pro-domain affects the flexibility in the core alpha helix of each protein in opposite ways. As mentioned above, in ISP1, the presence of the pro-domain rigidifies this core helix as seen by the increase in fluctuations in the absence of the pro-domain as measured by hydrogen exchange. Thus, the increase in stability provided by the pro-region is transmitted to the core alpha helix and appears to rigidify this region. We see the opposite trend in the extracellular protein SbtE, where the presence of the pro-domain increases the flexibility of the core helix. In fact, we see no measurable hydrogen exchange for this helix in the mature protein. Interestingly, while the relatively large pro-domain dramatically increases the global stability of the protein, it generates increased flexibility within the core. How these energetic effects are uncoupled is unclear.

### Sequence basis for the observed differences in the energy landscape

While ISP1 and SbtE have similar folds, they have only 45% sequence identity – making it difficult to identify the sequence basis for the differences in their energy landscape. The observed differences in the flexibility of the core alpha-helix suggest a potential region of the protein to focus on. In fact, ISP1 has an insertion at the C-terminus of the alpha helix that is conserved amongst many ISPs (Fig. 8). This insertion may be one source of the difference in folding/unfolding mechanisms between SbtE and ISP1. Furthermore, the active site serine resides at the N-terminus of this helix and in the case of ISP1, this whole region becomes more flexible in the absence of the pro-domain. Increased dynamics may benefit the enzymatic ability of ISP1. Since ISP1 does not reside in harsh extracellular conditions there is no reason to maintain a rigid core, and therefore a rigid native state. How SbtE maintains activity with such a rigid core is unclear.

**Figure 8:**
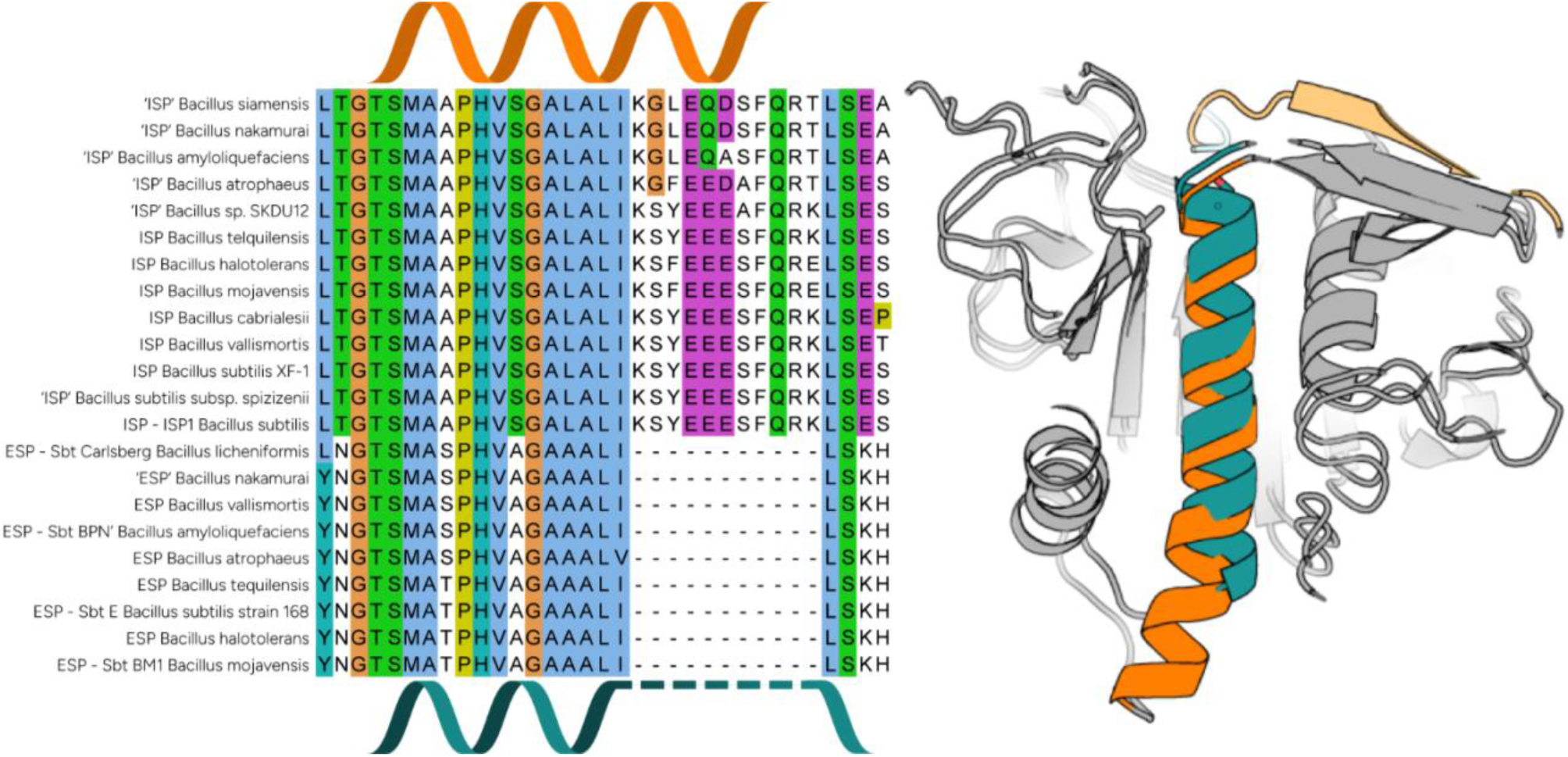
ISPs have a conserved insertion at the C-terminus of the core alpha helix. Left – Section of sequence alignment showing core alpha helix sequence from ISPs (top of alignment) and ESPs (bottom of alignment) from Bacillus species. ISP core alpha helix region is highlighted by orange ribbon cartoon and ESP core alpha helix region is highlighted by blue ribbon cartoon. Right – Slice of structural alignment of ISP1 and SbtE showing elongated core alpha helix in ISP1 (orange, AlphaFold3 model) versus the core alpha helix in SbtE (blue, PDB: 1SCJ). A histogram of each sequences pro-domain length can be found in Figure S7.

### Comparison between SbtE and alpha-lytic protease

A widely accepted hypothesis for the evolution of kinetically stable secreted proteases is that, once secreted into the extracellular space, they need to avoid degradation by the many other proteases present in the environment. For most proteins, proteolysis requires fluctuation from the native conformation (Park & Marqusee, 2004). Indeed, αLP was shown to have extremely limited fluctuations from the native state over the course of six months and outlast trypsin and chymotrypsin in an *in vitro* competition (Jaswal et al., 2002a). Over half of αLP’s protons were found to have a protection factor (P_f_ = k_int_/k_obs_) of greater than 10^4^ and 18 protons with a P_f_ of greater than 10^9^. While HDX-MS does not allow determination of residue-specific protection factors (due to measurements at the peptide vs amide level), the estimated P_f_’s for SbtE range from ∼10^0^ to 10^6^, with only a few greater than 10^4^.

Surprisingly, our results suggest that such complete suppression of fluctuations is not needed for increased proteolytic resistance, since SbtE does show equilibrium fluctuations, albeit rare, and nevertheless seems to outcompete thermolysin in a competition assay. Thus, while SbtE may have a greater degree of flexibility than αLP, it is still quite resistant to proteolysis. Additionally, ISP1 demonstrates similar proteolytic resistance, albeit through a different mechanism whereby ISP1 maintains a significant folded (uncleavable) population due to thermodynamics. Perhaps there exist different selective pressures in different environments requiring different levels of rigidity – or the increased resistance to proteolysis of αLP is not the only evolutionary pressure on the rigidity of this protein.

In sum, we have examined two homologous proteins from the subtiliase family with similar folds and sequences. We have found that despite these similarities the two proteases have very different energetics at the global and local levels. Differences in their local energetics within the core alpha helix have revealed that an insertion into this core helix may be an evolutionarily important feature of their landscape. How these features and kinetic barriers are encoded in protein sequence remains an outstanding question in the field. Our study suggests the need for high throughput experiments to explore the sequence dependence of a protein’s kinetic barriers for a specific fold.

## Materials and Methods

### Plasmid Cloning

The DNA encoding intracellular and extracellular subtilisin were purchased as gBlock gene fragments from IDT. Enzyme digestion and site-directed mutagenesis were used to clone: (1) Pro-Subtilisin E S221A and Pro-Subtilisin E WT into a pSV272 expression vector without the secretion signal and an N-terminal His6-MBP-TEV-(GS)25-PreScission tag and (2) Pro-ISP1 S250A and ISP1 S250A into pET-27b(+) (Novagen) expression vector. Sequences were confirmed with Sanger sequencing.

### Extracellular Subtilisin Protein Expression

Bl21 (DE3)pLysS competent cells were transformed with pSV272 vectors containing either Pro-SbtE S221A or Pro-SbtE WT DNA and plated on LB agar plates with 50 μg/mL kanamycin. Starter cultures were inoculated with single colonies for overnight growth to saturation. For Pro-SbtE S221A, 15 mL of LB media overnight culture was used to inoculate 1 L of LB media with 50 μg/mL kanamycin. Cells were grown at 37°C and 250 rpm to an optical density at 600 nm of 0.6-0.8 and induced with 1 mM IPTG. Cells were harvested after 3 hrs by centrifugation at 4000 g for 30 min at 4°C. For Pro-SbtE WT, 100 μL of 2xYT media overnight culture was used to inoculate 5 mL of 2xYT media with 50 μg/mL kanamycin and 1% glucose and grown at 37°C and 250 rpm to an optical density at 600 nm of 0.6-0.8. Cells were cooled to 20°C and induced with 1 mM IPTG. Cells were harvested after 18 hrs by centrifugation at 4000 g for 10 min at 4°C. Cell pellets were stored at −80°C until purification.

### Intracellular Subtilisin Protein Expression

Bl21 (DE3)pLysS competent cells were transformed with pET-27b(+) vectors containing either Pro-ISP1 S250A or ISP1 S250A DNA and plated on LB agar plates with 50 μg/mL kanamycin. Starter cultures were inoculated with single colonies for overnight growth to saturation. For Pro-ISP1 S250A and ISP1 S250A, 15 mL of LB media overnight culture was used to inoculate 1 L of LB media with 50 μg/mL kanamycin. Cells were grown at 37°C and 250 rpm to an optical density at 600 nm of 0.6-0.8 and induced with 1 mM IPTG. Cells were harvested after 3 hrs by centrifugation at 4000 g for 30 min at 4°C.

### Pro-Subtilisin E S221A Protein Purification

Pellets were thawed on ice, resuspended in 30 mL buffer A1 (50 mM Tris pH 8, 150 mM NaCl, 1 mM CaCl_2_, 0.5 mM TCEP) with 1 μL benzonase per 1 L pellet, and lysed by sonication at 30% for 15 minutes on ice. The lysate was centrifuged at 13000 rpm for 30 min at 4°C. The resulting pellet was washed once with 30 mL of 1% Triton in buffer A1 and once with 30 mL of buffer A1 alone. The washed inclusion bodies were resuspended in 10 mL of buffer A with 8 M urea, stirred at RT for 15 minutes, and sonicated for 10 seconds 5x. The denatured inclusion bodies were centrifuged at 14000g for 15 minutes. Protein was refolded by dropwise 10-fold dilution into buffer A1, stirred at RT for 15 minutes, and centrifuged at 13000 rpm for 30 minutes.

The supernatant was incubated with amylose beads for 2 hrs rotating at 4°C. The amylose beads were loaded into a gravity column. Flow through was collected and beads were washed with 50 mL of buffer A. His6-MBP-TEV-(GS)25-PreScission-Pro-SbtE S221A was eluted from the amylose beads in three 10 mL fractions with 10 mM maltose in buffer A1. The protein concentration of each fraction was measured by ultraviolet-visible (UV-vis) spectroscopy, fractions with protein were pooled, and PreScission protease was added, and the cleavage reaction proceeded overnight at 4°C. The N-terminal His6-MBP-TEV-(GS)25-PreScission tag was removed with subtractive a NiNTA step. The cleavage reaction was diluted to 50 mL in buffer A1 to 2.5 mM imidazole and incubated with NiNTA beads for 2 hrs rotating at 4°C. The NiNTA beads were loaded into a gravity column. Flow through was collected and beads were washed with 50 mL of buffer A1 with 10 mM imidazole. Flow through and wash fractions were both concentrated to <1 mL using Amicon Ultra-15 Centrifugal Filter with a 10 kDa MWCO. The protein concentration was measured by UV-vis spectroscopy and fractions with protein were pooled. The pooled fractions were 0.22 μm filtered and further purified by size-exclusion chromatography on a HiLoad S75 16/600 or HiLoad S200 16/60 (GE) in buffer B (50 mM Tris pH 8, 500 mM (NH_4_)_2_SO_4_, 1 mM CaCl_2_, 0.5 mM TCEP). The peak corresponding to Pro-SbtE S221A was collected, concentration was measured by UV-vis spectroscopy and stored overnight at 4°C for immediate use.

### Active Subtilisin E Protein Purification

Pellets were thawed on ice, resuspended in 0.5 mL BugBuster (Millipore) with 0.5 μL benzonase per 5 mL pellet, and rotated at RT for 2 hrs Lysis released soluble, active SbtE, which cleared the majority of the lysate. Lysate was centrifuged at 14000g for 10 minutes, concentrated to <2 mL, and 0.22 μm filtered. Filtered supernatant was run on a HiLoad S75 16/600 or HiLoad S200 16/60 (GE) in buffer B. The peak corresponding to SbtE was collected and concentrated to <2 mL. SbtE was diluted 10-fold in buffer A2 (50 mM Tris pH 8, 25 mM NaCl, 1 mM CaCl_2_, 0.5 mM TCEP) and further purified on a HiTrap Q HP (GE) and SbtE was collected from the flow-through. The peak corresponding to SbtE was collected, concentration was measured by UV-vis spectroscopy, and stored at 4°C.

### Pro-ISP1 S250A Protein Purification

Pellets were thawed on ice, resuspended in buffer A2 with 1 μL benzonase per 1 L pellet, and lysed by sonication at 30% for 15 minutes on ice. The lysate was centrifuged at 13000 rpm for 30 min at 4°C. Supernatant was 0.22 μm filtered. Pro-ISP1 S250A was isolated using anion-exchange chromatography on a HiPrep 16/10 Q XL (GE) and eluted with a gradient from 25 mM to 1 M NaCl. Pro-ISP1 S250A typically elutes around 300 mM NaCl. The peak corresponding to Pro-ISP1 S250A was collected, concentrated to <2 mL, and 0.22 μm filtered. Pro-ISP1 was further purified by size-exclusion chromatography on a HiLoad S75 16/600 or HiLoad S200 16/60 (GE) in buffer B. The peak corresponding to Pro-ISP1 S250A was collected, concentration was measured by UV-vis spectroscopy and stored overnight at −80°C for later use.

### ISP1 S250A Protein Purification

Pellets were thawed on ice, resuspended in buffer A2 with 1 μL benzonase per 1 L pellet, and lysed by sonication at 30% for 15 minutes on ice. The lysate was centrifuged at 13000 rpm for 30 min at 4°C. The resulting pellet was washed once with 30 mL of 1% Triton in buffer A2 and once with 30 mL of buffer A2 alone. The washed inclusion bodies were resuspended in 10 mL of buffer A2 with 8 M urea, stirred at RT for 15 minutes, and sonicated for 10 seconds 5x. The denatured inclusion bodies were centrifuged at 14000g for 15 minutes. Protein was refolded by dropwise 10-fold dilution into buffer A2, stirred at RT for 15 minutes, and centrifuged at 13000 rpm for 30 minutes. Supernatant was 0.22 μm filtered. ISP1 S250A was isolated using anion-exchange chromatography on a HiPrep 16/10 Q XL (GE) and eluted with a gradient from 25 mM to 1 M NaCl. ISP1 S250A typically elutes around 250 mM NaCl. The peak corresponding to ISP1 S250A was collected, concentrated to <2 mL, and 0.22 μm filtered. Pro-ISP1 was further purified by size-exclusion chromatography on a HiLoad S75 16/600 or HiLoad S200 16/60 (GE) in buffer B. The peak corresponding to ISP1 S250A was collected, concentration was measured by UV-vis spectroscopy and stored overnight at 4°C for immediate use.

### Equilibrium Denaturation by Circular Dichroism Spectroscopy

Protein samples were diluted to a concentration of 0.04 mg/mL in 0 M or 7 M GdmCl in buffer B. High and low denaturant protein samples were either mixed manually to create a range of GdmCl concentrations or high denaturant protein was titrated into low denaturant protein with the ATS-350 titrator (Jasco). Protein was allowed to equilibrate at each GdmCl concentration for at least 5 minutes at 25°C before measurement. CD signal was measured at 222 nm for 30 seconds in a 1 cm quartz cuvette (Starna) with a stirring magnetic stir bar. For manually mixed samples, each sample was recovered and the GdmCl concentration was measured with a refractometer. Denaturation curves were plotted and fit to either a standard six parameter two state fit (Eq. 2) or a nine parameter three state fit (Eq. 3) (Barrick & Baldwin, 1993; Street, Courtemanche, & Barrick, 2008).

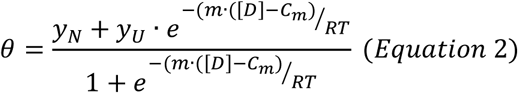

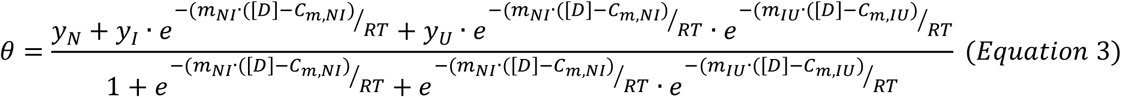

Where Θ is the CD signal at 222 nm, m is the m-value for each transition, and:

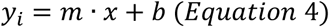

### Unfolding Kinetics by Circular Dichroism Spectroscopy

Protein samples were diluted to a concentration of 0.4 mg/mL in 0 M GdmCl in buffer B. All samples and buffers were allowed to equilibrate overnight at 25°C. For each experiment 1.95 mL of unfolding buffer was placed into a 1 cm quartz cuvette. To initiate unfolding, 0.05 mL of protein was added to the same cuvette, rapidly mixed with p1000 micropipetter, capped, and placed in the CD with a stirring magnetic stir bar. An estimate of the deadtime for mixing and starting the measurement was also recorded. CD signal was measured at 222 nm, integrated over 8 seconds, every 10 seconds for 3 to 8 hours. Each sample was recovered and the final GdmCl concentration was determined with a refractometer. Individual unfolding traces were plotted and fit to a single exponential (Eq. 5). Calculated unfolding rates were plotted against GdmCl concentration and fit to a line (Eq. 4) in Python.

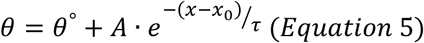

Where Θ is the CD signal and τ is the lifetime of the starting state.

### Unfolding and Refolding Kinetics by Stopped-Flow Fluorescence

Protein sample was diluted to a concentration of 150 μM in 0 M or 7 M GdmCl in buffer B. All samples and buffers were allowed to equilibrate for at least 5 minutes at 25°C. Protein sample was loaded into 2 mL slip tip syringe and attached to position 4 on the SFM 400 (BioLogic) and buffer was loaded into a 10 mL slip tip syringe and attached to position 3. Positions 1 and 2 were loaded with MilliQ water and not used during the experiment. For each shot of a refolding experiment 225 μL of buffer and 25 μL of protein were pushed into the FC-15 cuvette (BioLogic) at a flow of 6.5 mL/s. Five shots were used to clear the cuvette and the following five shots were recorded. Sample was excited at 280 nm and emission at 320 nm was recorded for 50-500 ms sampling every 0.01-1 ms. The dead time of the stopped-flow was 5.6 ms. Individual unfolding traces were plotted and fit with Python scripts with a single exponential equation (Eq. 5). Calculated unfolding rates were plotted against GdmCl concentration and fit to a line (Eq. 4) in Python.

### Unfolding and Refolding Kinetics by Stopped-Flow Circular Dichroism Spectroscopy

Protein sample was diluted to a concentration of 7 mg/mL in 0 M or 7 M GdmCl in buffer B. All samples and buffers were allowed to equilibrate for at least 5 minutes at 25°C. Protein sample was loaded into reservoir 1 of the Aviv 430 stopped flow and buffer was loaded into reservoir 3. For each shot of a refolding experiment 80 μL of buffer and 8 μL of protein were pushed into 0.1-cm path length cuvette using a delta mixer (Aviv). Five shots were used to clear the cuvette, and the following five shots were recorded. Signal at 222 nm was recorded for 50-500 ms sampling every 0.01-1 ms. The stopped-flow dead time was 70 ms. Individual unfolding traces were plotted and fit with Python scripts with a single exponential equation (Eq. 5). Calculated unfolding rates were plotted against GdmCl concentration and fit to a line (Eq. 4) in Python.

### Subtilisin E Activity Standard Curve

SbtE was serially diluted to a concentration of 2×10^1^to 2×10^−5^ μM in 80% buffer B, 20% buffer C. The substrate N-succinyl-Ala-Ala-Pro-Phe-P-nitroanilide (s-AAPF-pNA) was prepared by dilution from N,N-dimethylformamide (DMF) into buffer B and its concentration was measured by absorption at 315 nm. Phenylboronic acid was dissolved in buffer B. The enzymatic reaction was initiated by adding 6 μL of Subtilisin E standard to 54 μL of in buffer B for a final concentration of 1 μM substrate, 7.5 μM phenylboronic acid in a 1 cm masked quartz cuvette (Starna). This was done for all Subtilisin E standards, measuring activity of 2 to 2×10^−6^ μM protein at 25°C. Absorption at 410 nm was recorded every 100 ms for 10 minutes. Individual activity traces were plotted and fit to a line (Eq. 4). The detection limit is approximately 2 pm in 60 μL.

### Refolding Kinetics Measured by Subtilisin E Activity

This protocol was adapted from (Sohl et al., 1998). To determine the refolding rate, at timepoints 24, 48, 72, and 168 hours both protease activity arising from refolded [N], and the decrease in total protein [I] attributed to autodigestion by refolded N were measured as described below. Fraction folded ([*N*]⁄[*I*]*_t_*) was plotted against time and fit with *k_f_* = ([*N*]⁄[*I*]_0_)⁄t (Sohl et al., 1998) to find the estimated refolding rate.

Protein was first unfolded by dialyzing into 7 M GdmCl in buffer C (50 mM Potassium Acetate pH 5, 500 mM (NH_4_)_2_SO_4_, 1 mM CaCl_2_, 0.5 mM TCEP) with 10 mM phenylboronic acid, a low-affinity (K_i_ = 8 mM or 67 μM – (Keller, Seufer-Wasserthal, & Jones, 1991; Lindquist & Terry, 1974) subtilisin inhibitor, overnight at 4°C. This procedure limits autodigestion and removes any protein fragments from residual autodigestion during unfolding. The unfolding incubation time is several orders of magnitude longer than the measured τ_folded_ at 7 M GdmCl to allow for complete unfolding. Refolding was initiated by rapidly diluting 5 μL of 400 μM protein with 995 μL of buffer C without subtilisin inhibitor (leading to a diluted inhibitor concentration of 75 μM) at 4°C or 25°C.

To determine the concentration of [N], 6 μL of refolding reaction to 54 μL s-AAPF-pNA in buffer B for a final concentration of 0.2 μM protein, 1 μM substrate, and 7.5 μM phenylboronic acid, and enzyme activity was monitored as described for the SbtE standard curve. Because not activity was detected, the maximum possible [N] refolded at each timepoint was estimated as less than or equal to the detection limit of the assay (2 pM in the 60 μL reaction) determined from the SbtE Activity Standard Curve. The decrease in intact folding intermediate (I or total protein) due to proteolysis by refolded N was monitored by first adding 2 molar excess of the pro-domain to 6 μL of the refolding reaction and equilibrating at 25°C for 2 hours to enable pro-catalyzed refolding of undigested [I]. The sample was then diluted 5-fold with buffer B and treated with 0.004 μg of trypsin and incubated at 25°C for 30 minutes to digest the pro-domain. The enzymatic reaction was initiated by addition of substrate to 1 μM s-AAPF-pNA in 60 μL of the treated sample and monitored as described.

### Protease Survival Assay

Stock thermolysin was prepared by dissolving ∼0.5 mgs of thermolysin (Sigma-Aldrich, P1512) into buffer B and concentration was determined by measuring absorption at 280 nm. Proteolysis was initiated by adding the stock thermolysin to Subtilisin E in buffer B for a final concentration of 10 μM and 10 μM respectively and incubated at 25°C. Samples were removed at at 0.5, 1, 2, 4, 8, 24, 48, 72, 96, 168, and 336 hours and quenched with 100 mM EDTA or 100 mM EDTA and 250 μM PMSF. Each time point was run on a NuPAGE Bis-Tris gel (Invitrogen), stained with Sypro Tangerine, and imaged on a Typhoon imager. Band intensities were quantified in ImageJ. Data was plotted and fit to a single exponential equation.

### Continuous Hydrogen-Deuterium Exchange

For all hydrogen-deuterium exchange (HDX) experiments, buffer B was lyophilized and resuspended in D_2_O. Deuterated buffer B and protein samples were pre-equilibrated to 25°C. HDX was initiated by diluting protein samples 10-fold into deuterated buffer B for a final protein concentration of 5 μM. At each time point, samples were quenched by mixing 50 μL of HDX reaction with 50 μL of ice-cold 2x quench buffer (3.5 M GdmCl, 1.5 M Glycine pH 2.4, 500 mM TCEP) and flash freezing in liquid nitrogen. Samples were stored at −80°C. For Pro-SbtE S221A, Pro-ISP1 S250A, and ISP1 S250A, 0.5 μg/mL angiotensin peptide was included as a fiduciary as a back-exchange control, although the data were not back-exchange corrected.

### Protease Digestion and LC-MS

Protein samples were injected into the LC (Thermo Ultimate 3000) and run on the Q Exactive Orbitrap Mass Spectrometer (ThermoFisher) as described previously (Costello et al., 2022). The quenched samples were inline digested with columns made by covalently attaching aspergillopepsin (Sigma-Aldrich, P2143) and the porcine pepsin (Sigma-Aldrich, P6887) to resin using standard aldehyde chemistry. Digestion occurred at a flow rate of 200 μL/min of aqueous phase buffer (0.1% formic acid) at 10C. Peptides were then desalted for 4 minutes on a hand-packed trap column (Thermo Scientific POROS R2 reversed-phase resin 1112906, IDEXC-130B). Peptides were separated on a C18 analytical column (Waters Acquity UPLC BEH C18 Column, 1.7 μm particle size, 1 mm (inner diameter) × 50 mm, 186002344) with a gradient 5-40% organic phase buffer (100% acetonitrile, 0.1% formic acid) at 40 μL/min over 14 minutes. The analytical and trap columns were then washed with 40-90% organic phase buffer sawtooths every 30 seconds. The analytical and trap columns were then equilibrated at 5% organic phase buffer before the next injection. After digestion and desalting, the protease columns were washed with three injections of 100 μL of 1.6 M GdmCl, 0.1% formic acid before the next injection. Peptides were eluted directly into Q Exactive Orbitrap Mass Spectrometer operating in positive mode (resolution 70000, AGC target 3 × 106, maximum IT 50 ms, scan range 300–1,500 m/z). For each protein variant, a tandem mass spectrometry experiment was performed (full MS settings were the same as above, and data-dependent MS2 settings included 17,500, automatic gain control target of 2 × 105, maximum injection time of 100 ms, loop count of 10, isolation window of 2.0 m/z, normalized collision energy of 28, charge state of 1 and ≥7 excluded and dynamic exclusion of 15 s) on undeuterated samples. LC and MS methods were run using Xcalibur 4.1 (Thermo Scientific).

### Peptide Identification

Peptides were identified from tandem MS data using Byonic (Protein Metrics) as described previously (Costello et al., 2022). The sequence of the expressed construct with the angiotensin peptide sequence added to the C-terminal of the sequence, if present, was used as the search reference. The search parameters were set as follows: specificity to nonspecific, precursor mass tolerance to 6 ppm, and fragment mass tolerance to 20 ppm.

### HDExaminer 3 Analysis

Peptide isotope distributions at each time point were fit in HDExaminer 3 as previously described (Costello et al. 2022). Peptide isotope distributions for each time point were fit in HDExaminer 3. Deuteration levels were determined by subtracting the mass centroids of deuterated peptides from undeuterated control peptides. All analyzed peptides showed EX2 behavior or no exchange.

## Supporting information

FullAlignment

ISP1_SbtE_Alignment

## Acknowledgements

We would like to thank Sophie Shoemaker for help with HDX-MS, Eva Gerber for technical advice, the entire Marqusee Lab for advice and suggestions, and David Agard and his lab for advice and reagents. This work was funded by a grant from the NIH (GM 149319). SM is a Chan Zuckerberg Biohub Investigator.

## Abbreviations and symbols

αLP: alpha-lytic proteases
CD: circular dichroism
DMF: dimethylformamide
ESP: extracellular subtilisin protease
GdmCl: guanidinium chloride
HDX: hydrogen deuterium exchange
ISP: intracellular subtilisin protease
ISP1: intracellular subtilisin protease 1
LC: liquid chromatography
MS: mass spectrometry
Pro-ISP1: pro-intracellular subtilisin protease 1
Pro-SbtE: pro-subtilisin E
SbtE: subtilisin E
SF: stopped flow
UV-vis: ultraviolet-visible

