## Supplement for "Exploring the sequence and structural determinants of the energy landscape from thermodynamically stable and kinetically trapped subtilisins: ISP1 and SbtE"

**This PDF file includes:**

Figs. S1 to S9

**Other supplementary materials for this manuscript include:**

ISP1_ProSbtE_muscle_I20240830-075152-0887-55872660-p1m.fasta
FullAlignment_muscle_I20240823-224802-0966-85644145-p1m.fasta

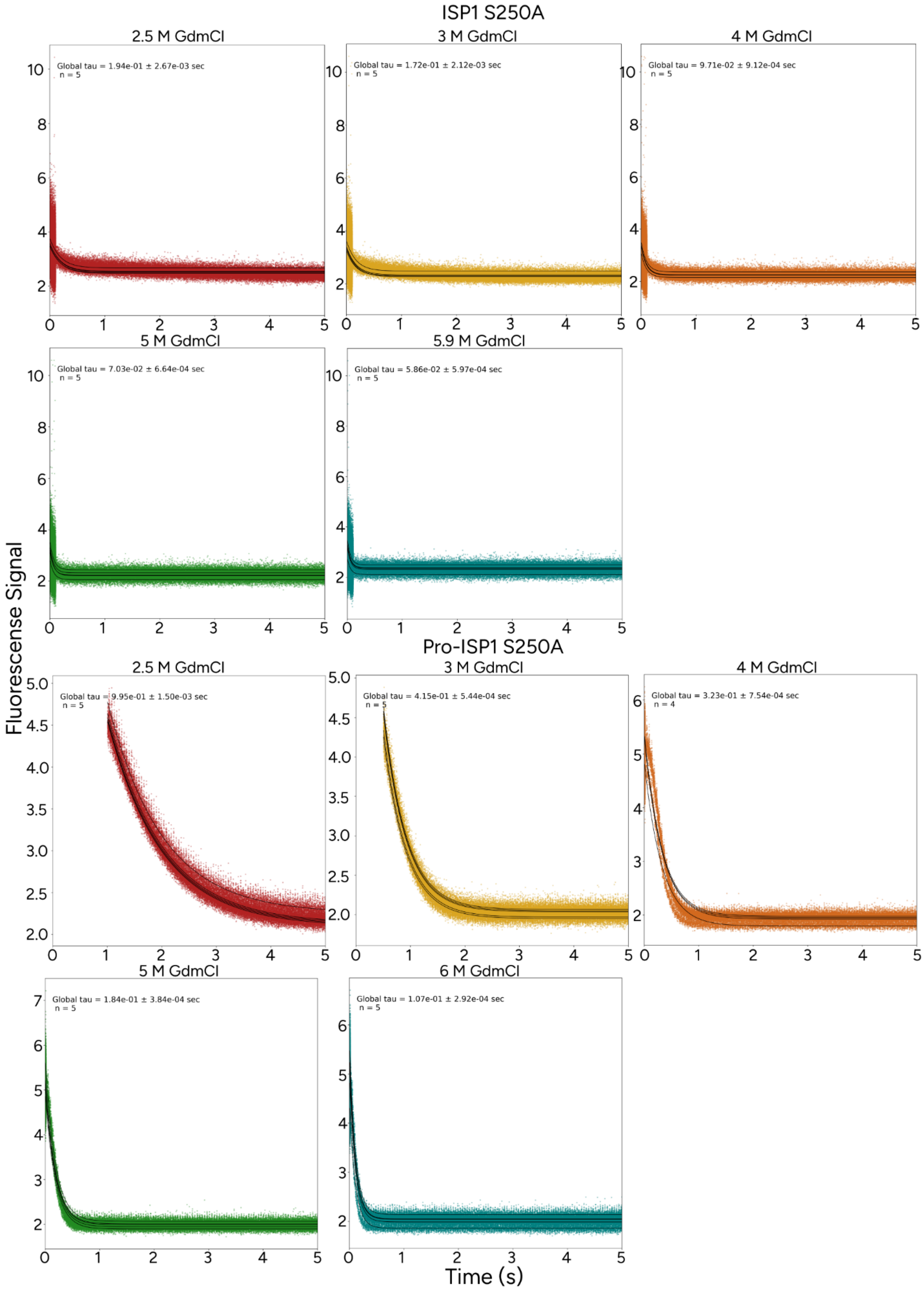
ISP1 S250A
2.5 M GdmCl
Global tau = 1.94e-01 ± 2.67e-03 sec
n = 5
3 M GdmCl
Global tau = 1.72e-01 ± 2.12e-03 sec
n = 5
4 M GdmCl
Global tau = 9.71e-02 ± 9.12e-04 sec
n = 5
5 M GdmCl
Global tau = 7.03e-02 ± 6.64e-04 sec
n = 5
5.9 M GdmCl
Global tau = 5.86e-02 ± 5.97e-04 sec
n = 5
Pro-ISP1 S250A
2.5 M GdmCl
Global tau = 9.95e-01 ± 1.50e-03 sec
n = 5
3 M GdmCl
Global tau = 4.15e-01 ± 5.44e-04 sec
n = 5
4 M GdmCl
Global tau = 3.23e-01 ± 7.54e-04 sec
n = 4
5 M GdmCl
Global tau = 1.84e-01 ± 3.84e-04 sec
n = 5
6 M GdmCl
Global tau = 1.07e-01 ± 2.92e-04 sec
n = 5
Fluorescense Signal
Time (s)

**Figure S1: Representative unfolding curves and single exponential fits from kinetic refolding studies of Pro-ISP1 S250A and ISP1 S250A (25°C, pH 8).** Pro-ISP1 S250A (left) and ISP1 S250A (right) unfolding curves using stopped-flow mixing monitored by fluorescence. Each plot shows cumulative data from 4-5 shots. Data are shown as closed circles. Fits for each replicate are shown as a solid line.

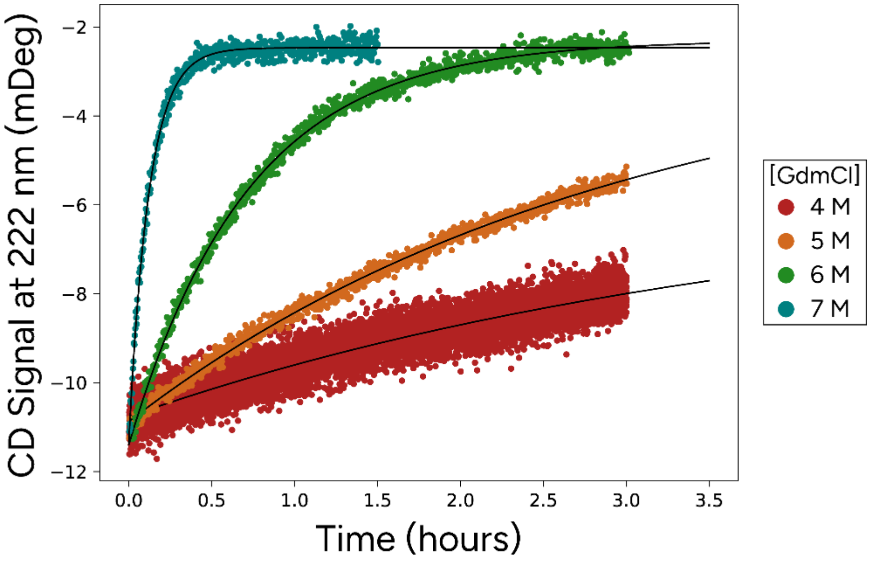


**Figure S2: Representative unfolding curves and single exponential fits from kinetic unfolding experiments on SbtE (25°C, pH 8).** Unfolding was initiated by manual mixing and monitored by CD at 222 nm. Each GdmCl concentration is color coded: 4 M in red, 5 M in orange, 6 M in green, 7 M in teal. Fits for each unfolding trace are shown as solid lines.

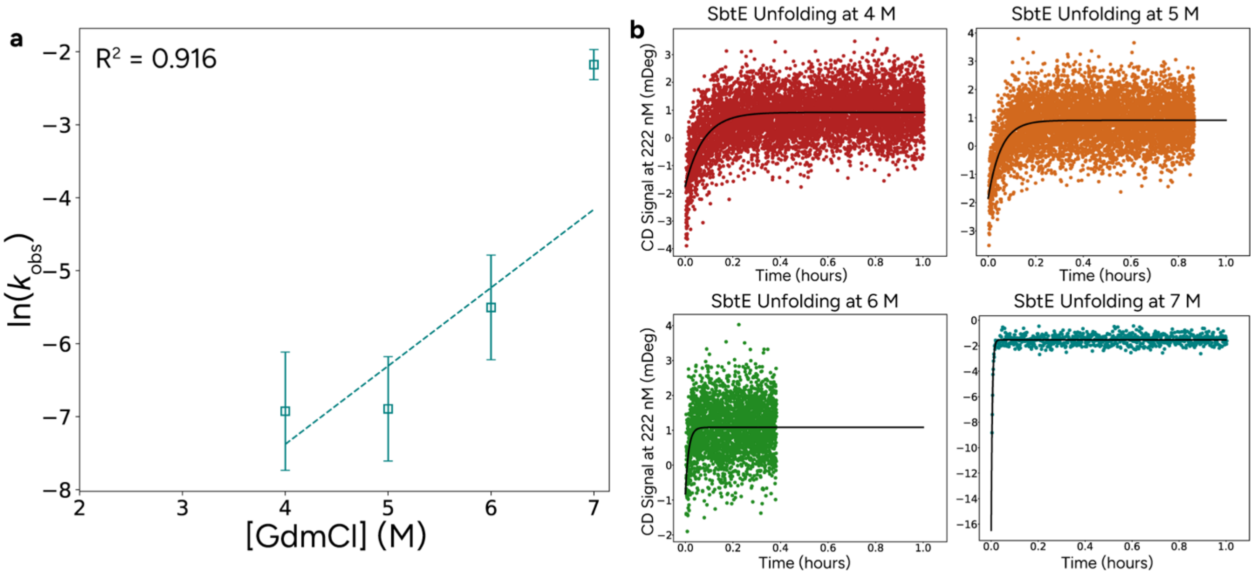


**Figure S3: Kinetic unfolding of SbtE (25°C, pH 5). a)** Unfolding as a function of guanidinium concentration for SbtE at pH 5 as monitored by CD at 222 nm. Data are shown as open squares with error bars showing standard error from three replicates. Linear regression is shown as dotted line. **b)** Representative unfolding curves and single exponential fits of SbtE unfolding by manual mixing monitored by CD at 222 nm. . Each GdmCl concentration is color coded: 4 M in red, 5 M in orange, 6 M in green, 7 M in teal. Fits for each unfolding trace are shown as solid lines.

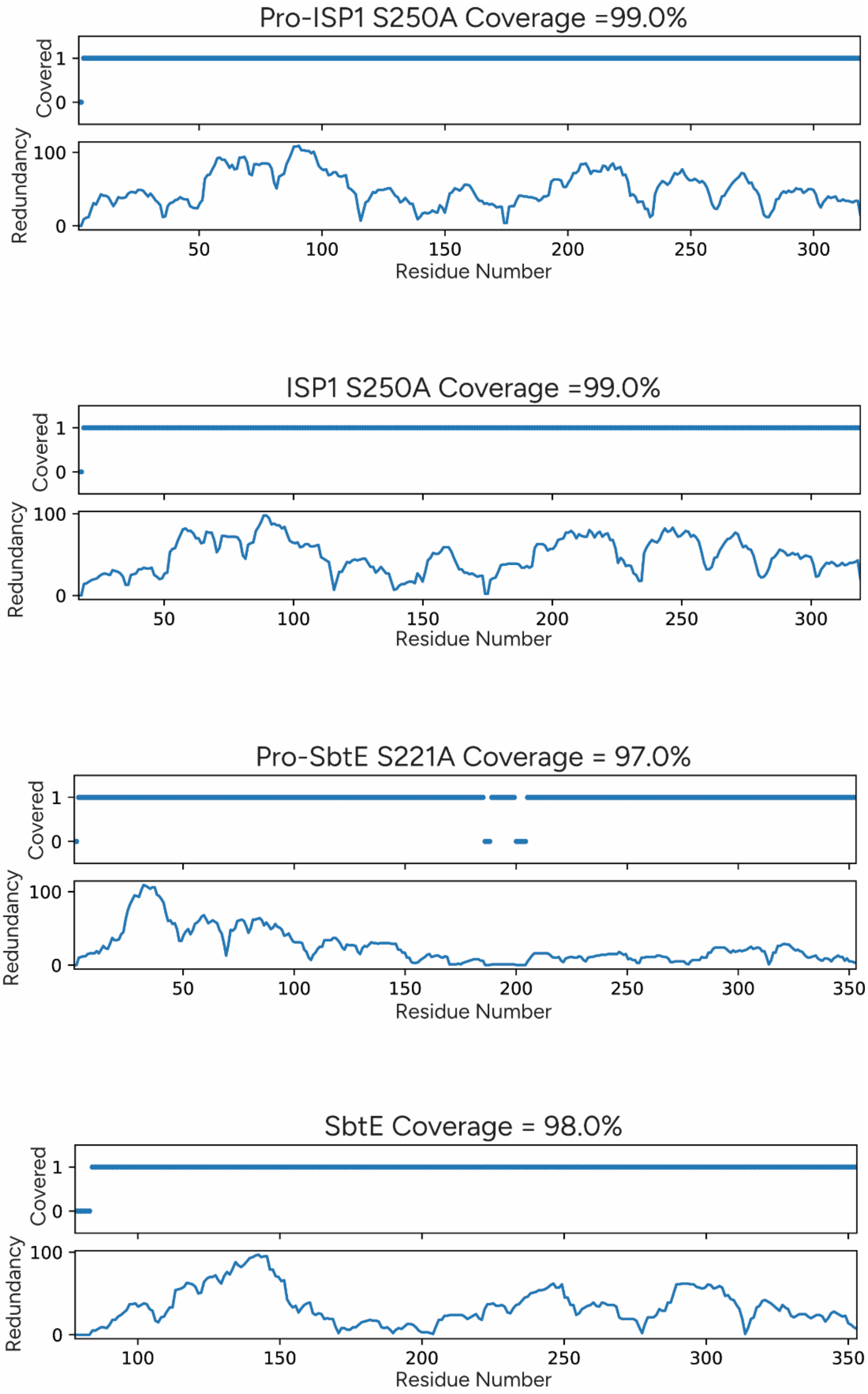


**Figure S4: Peptide coverage maps for Pro-ISP1 S250A, ISP1 S250A, Pro-SbtE S221A, and SbtE.** Peptide coverage and redundancy at each residue for all HDX experiments.

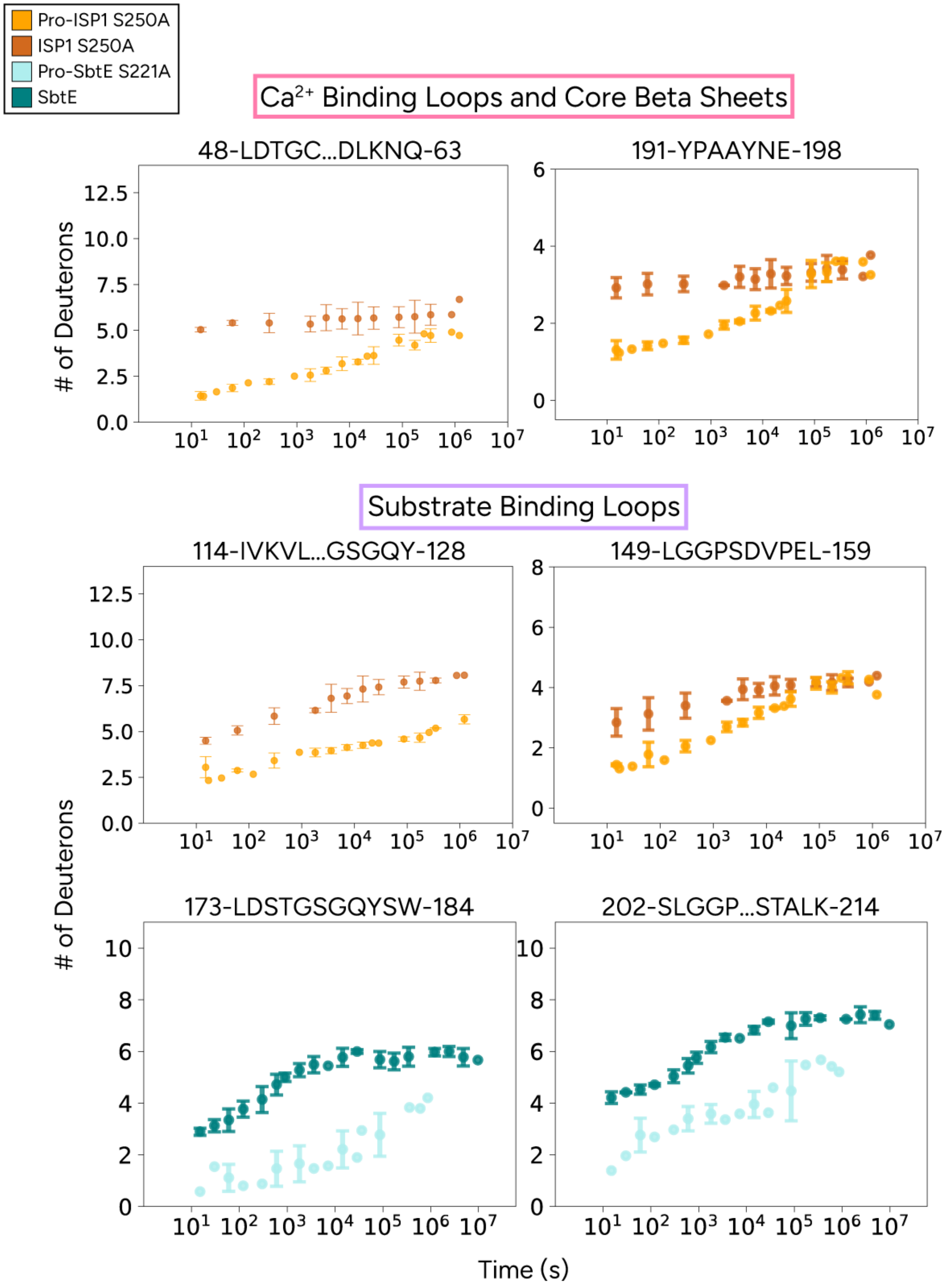


**Figure S5: Additional uptake plots for Ca2+ binding loops and core beta sheets in ISP1 and substrate binding loops in ISP1 and SbtE.** Uptake plots showing average number of deuterons per protein as a function of time for representative peptides (average of three replicates, error bars represent standard deviation not back exchange corrected). Plots are color coded by protein: Pro-ISP1 S250A in orange, ISP1 S250A in dark orange, Pro-SbtE S221A in light blue, SbtE in dark blue.

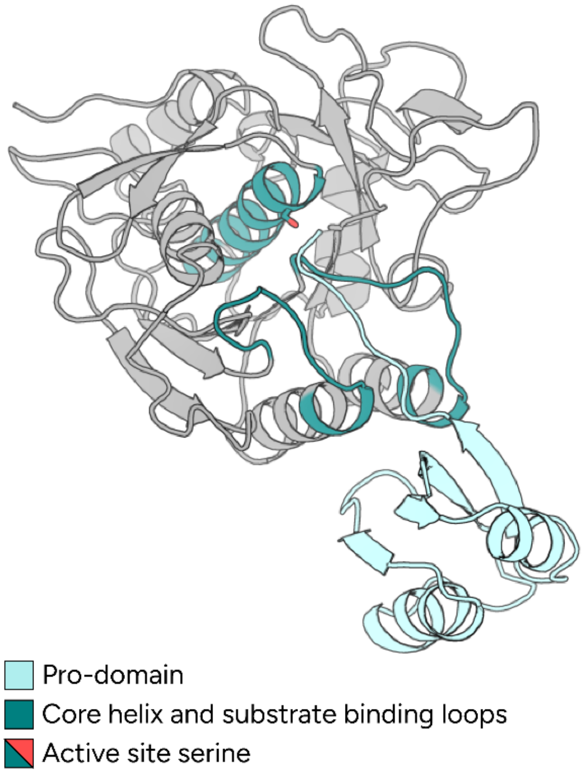


**Figure S6: Substrate binding loops become solvent exposed upon degradation/loss of pro-domain**

Ribbon structure of SbtE (PDB 1SCJ) with its pro-domain (left) and without its pro-domain (right). The pro-domain is shown in light blue. The core alpha helix and substrate binding loops are shown in teal. Active-site serine is shown as a stick in teal and red. The unstructured C-terminal end of the pro-domain sits between the substrate binding loops when the pro-domain is present.

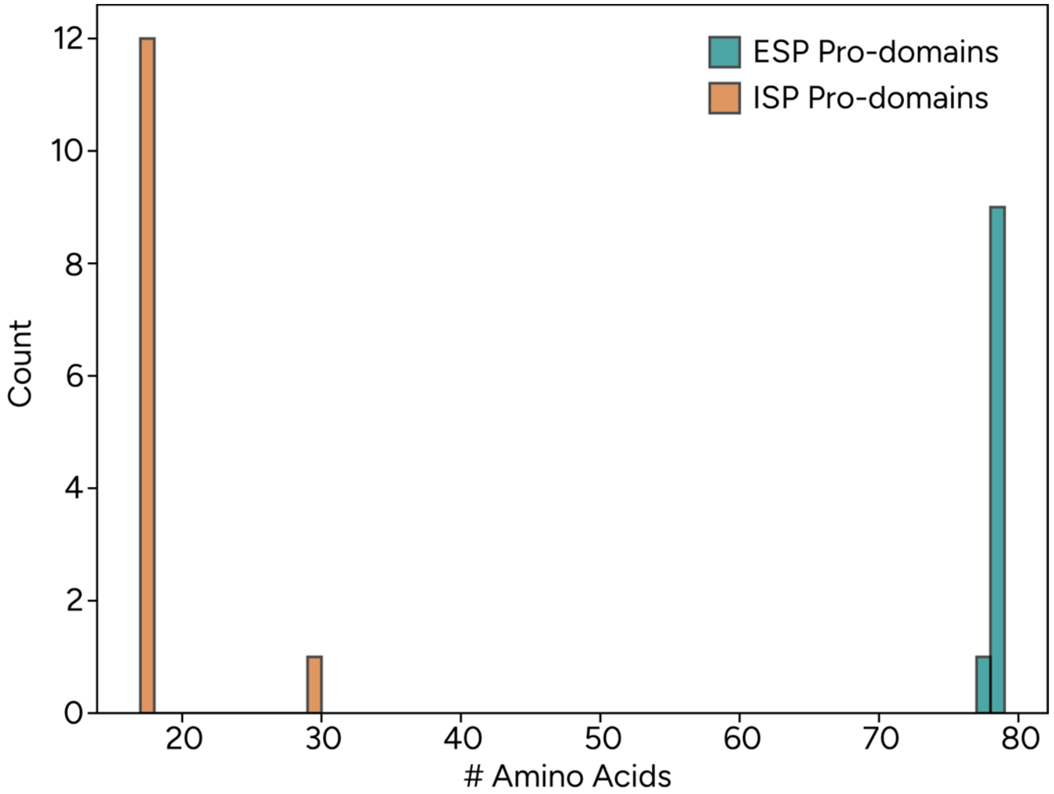


**Figure S7: Distribution of pro-domain size for aligned sequences from Figure 6** The number of each pro-domains of a given length are plotted as a distribution. Bars are color coded based on whether the pro-domains come from an ESP (blue) or an ISP (orange).

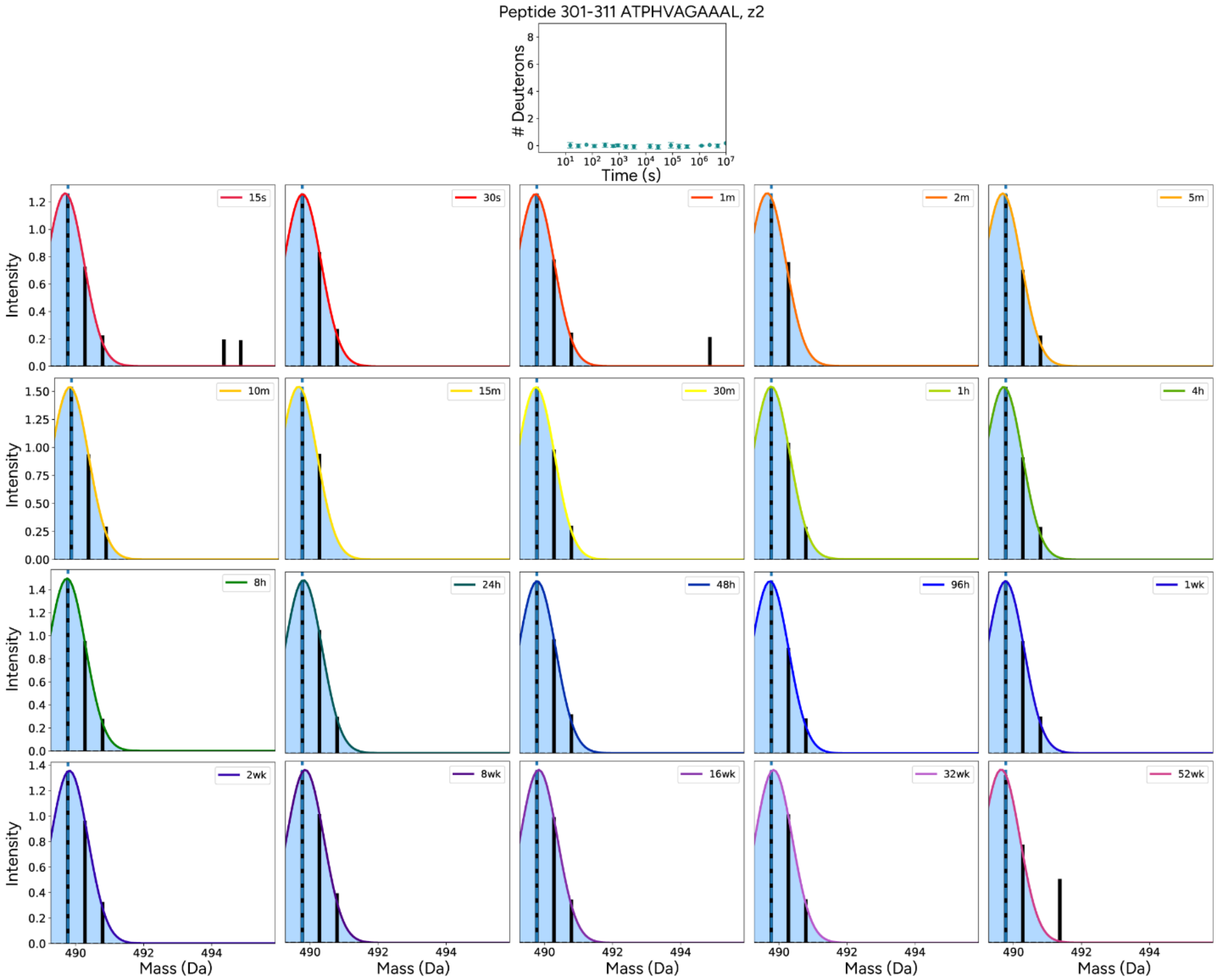


**Figure S8: Mass envelopes of peptide 301-311 does not shift over the course of a year.** An uptake plot of peptide 301-311 (top) shows the number of deuterons added per time point out to a year. Data are shown as closed circles and error bars represent standard deviation of three replicates. Mass spectra for each time point are shown as black bars with the first time point in the top left and last in the bottom right. Gaussian fits are shown as light blue and the centroid is marked by a dashed line.

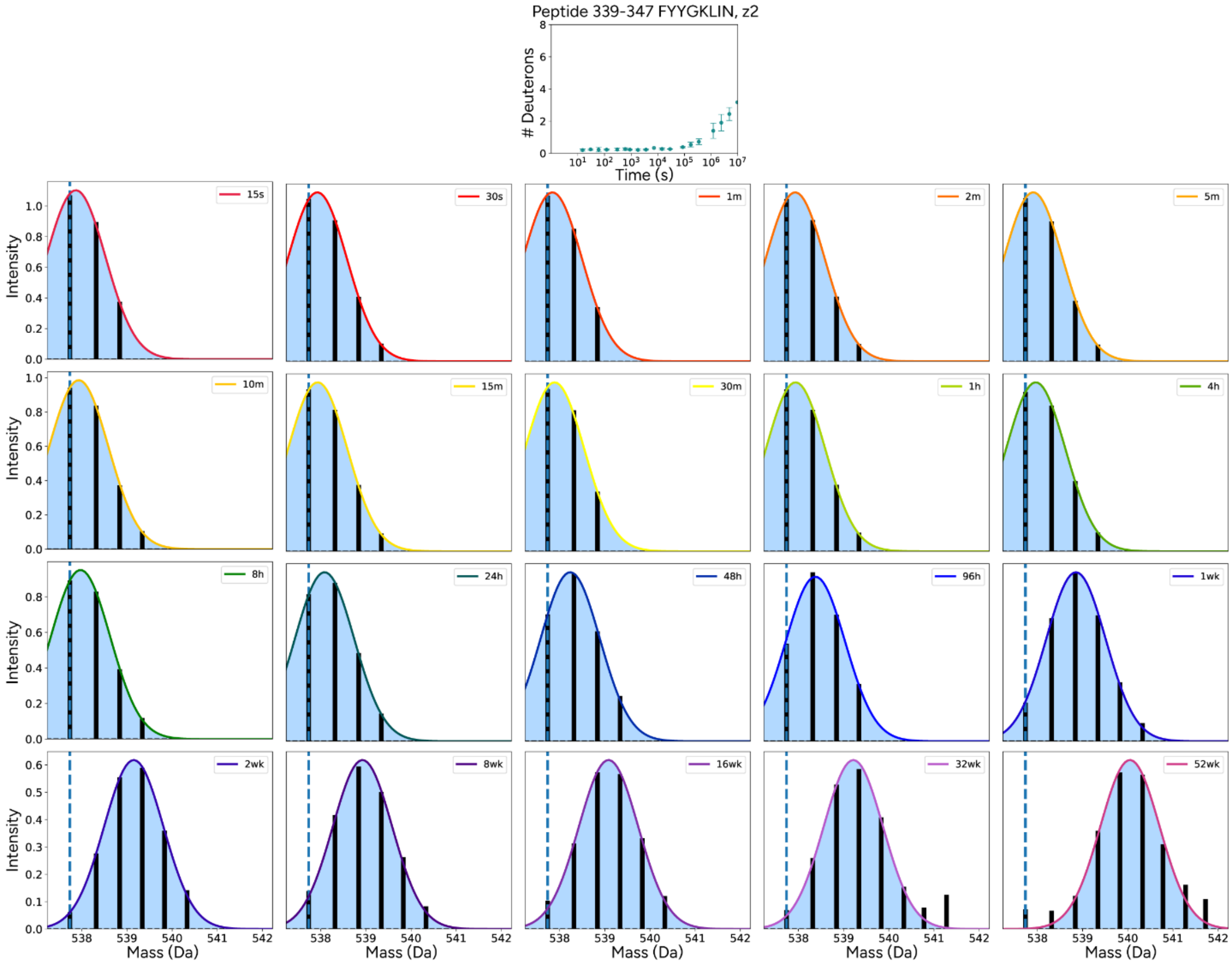


**Figure S9: Mass envelopes of peptide 339-347 (representative of core beta sheets) has a shifting centroid and no evidence of EX1 behavior.** An uptake plot of peptide 339-347 (top) shows the number of deuterons added per time point out to a year. Data are shown as closed circles and error bars represent standard deviation of three replicates. Mass spectra for each time point are shown as black bars with the first time point in the top left and last in the bottom right. Gaussian fits are shown as light blue and the centroid is marked by a dashed line.
